# Divergent Acyl Carrier Protein Decouples Mitochondrial Fe-S Cluster Biogenesis from Fatty Acid Synthesis in Malaria Parasites

**DOI:** 10.1101/2021.04.13.439690

**Authors:** Seyi Falekun, Jaime Sepulveda, Yasaman Jami-Alahmadi, Hahnbeom Park, James A. Wohlschlegel, Paul A. Sigala

## Abstract

*Plasmodium falciparum* malaria parasites are early-diverging eukaryotes with many unusual metabolic adaptations. Understanding these adaptations will give insight into parasite evolution and unveil new parasite-specific drug targets. Most eukaryotic cells retain a mitochondrial fatty acid synthesis (FASII) pathway whose acyl carrier protein (mACP) and 4-phosphopantetheine (Ppant) prosthetic group provide a soluble scaffold for acyl chain synthesis. In yeast and humans, mACP also functions to biochemically couple FASII activity to electron transport chain (ETC) assembly and Fe-S cluster biogenesis. In contrast to most eukaryotes, the *Plasmodium* mitochondrion lacks FASII enzymes yet curiously retains a divergent mACP lacking a Ppant group. We report that ligand-dependent knockdown of mACP is lethal to parasites, indicating an essential FASII-independent function. Decyl-ubiquinone rescues parasites temporarily from death, suggesting a dominant dysfunction of the mitochondrial ETC followed by broader cellular defects. Biochemical studies reveal that *Plasmodium* mACP binds and stabilizes the Isd11-Nfs1 complex required for Fe-S cluster biosynthesis, despite lacking the Ppant group required for this association in other eukaryotes, and knockdown of parasite mACP causes loss of both Nfs1 and the Rieske Fe-S protein in ETC Complex III. This work reveals that *Plasmodium* parasites have evolved to decouple mitochondrial Fe-S cluster biogenesis from FASII activity, and this adaptation is a shared metabolic feature of other *Apicomplexan* pathogens, including *Toxoplasma* and *Babesia*. This discovery also highlights the ancient, fundamental role of ACP in mitochondrial Fe-S cluster biogenesis and unveils an evolutionary driving force to retain this interaction with ACP independent of its eponymous function in FASII.

**Significance Statement:** *Plasmodium* malaria parasites are single-celled eukaryotes that evolved unusual metabolic adaptations. Parasites require a mitochondrion for blood-stage viability, but essential functions beyond the electron transport chain are sparsely understood. Unlike yeast and human cells, the *Plasmodium* mitochondrion lacks fatty acid synthesis enzymes but retains a divergent acyl carrier protein (mACP) incapable of tethering acyl groups. Nevertheless, mACP is essential for parasite viability by binding and stabilizing the core mitochondrial Fe-S cluster biogenesis complex via a divergent molecular interface lacking an acyl-pantetheine group that contrasts with other eukaryotes. This discovery unveils an essential metabolic adaptation in *Plasmodium* and other human parasites that decouples mitochondrial Fe-S cluster biogenesis from fatty acid synthesis and evolved at or near the emergence of *Apicomplexan* parasitism.

---

Malaria is an ancient scourge of humanity and remains a pressing global health challenge, especially in tropical Africa where hundreds of thousands of people die from malaria each year. *Plasmodium falciparum* malaria parasites are single-celled eukaryotes that evolved under unique selective pressures with unusual metabolic adaptations compared to human cells and well-studied model organisms such as yeast. Understanding the unique biochemical pathways that specialize parasites for growth within human red blood cells will shed light on their evolutionary divergence from other eukaryotes and unveil new parasite-specific targets for development of novel antimalarial therapies.

*P. falciparum* retains an essential mitochondrion required for biosynthesis of pyrimidines, acetyl-CoA, and Fe-S clusters (1-5). Although the parasite mitochondrion also contains enzymes involved in the citric acid (TCA) cycle and biosynthesis of ATP and heme, these pathways are dispensable for blood-stage parasites, which can scavenge host heme and obtain sufficient ATP from cytoplasmic glycolysis (6-10). In addition to these pathways, most eukaryotes, including mammals and yeast, also contain a mitochondrial type-II fatty acid biosynthesis (FASII) pathway that generates the octanoate precursor of the lipoic acid cofactor used by several mitochondrial dehydrogenases (11). In contrast to human and yeast cells, FASII enzymes in *P. falciparum* are absent from the mitochondrion and targeted instead to the apicoplast organelle (Table S1) (12). Although critical lipoate-dependent enzymes are present in the parasite mitochondrion (2), prior work has shown that these enzymes utilize scavenged lipoate obtained from the red blood cell rather than de novo synthesis (13).

The acyl carrier protein (ACP) is a key component of FASII and contains a strictly conserved Ser residue modified by a 4-phosphopantetheine (4-Ppant) group whose terminal thiol tethers the nascent acyl chain during fatty acid initiation, modification, and elongation (14). Consistent with FASII targeting to the apicoplast in *P. falciparum*, this organelle features a well-studied ACP homolog (PF3D7_0208500, aACP) that retains canonical ACP features, including the conserved Ser modified by a 4-Ppant group required for FASII function (12, 15). Although apicoplast FASII activity is essential for *P. falciparum* growth within mosquitoes and the human liver, this pathway is dispensable for blood-stage parasites which can scavenge fatty acids from the host (12, 16, 17).

Despite loss of mitochondrial FASII enzymes, *P. falciparum* has curiously retained a second ACP homolog (PF3D7_1208300) that is annotated as a mitochondrial ACP (mACP) but has not been directly studied. We noted that recent genome-wide knock-out studies in *P. berghei* and *P. falciparum* both reported that this gene was refractory to disruption, suggesting an essential, FASII-independent function (18, 19). We became interested to unravel what this essential function might be. Recent studies in yeast and human cells have identified an expanding network of mitochondrial ACP interactions beyond FASII that includes roles in respiratory chain assembly, Fe-S cluster biogenesis, and mitochondrial ribosomal translation (20-24). However, in all cases these critical interactions in yeast and/or humans are linked to mitochondrial FASII function and the Ppant group of ACP (23, 25-27).

We localized mACP to the *P. falciparum* mitochondrion and set out to understand its FASII-independent function in this organelle. Using conditional knockdown and immunoprecipitation studies, we discovered that mACP is essential for parasite viability and plays a critical role in binding and stabilizing the Isd11-Nfs1 cysteine desulfurase complex required for mitochondrial Fe-S cluster biogenesis. Unlike mACP interactions in yeast and humans, *P. falciparum* mACP binds to Isd11 via a divergent molecular interface that does not involve a 4-Ppant group. This work unveils a new molecular paradigm for essential mACP function without a Ppant group, underscores the critical and conserved role of mACP in mitochondrial Fe-S cluster biogenesis, and highlights a *Plasmodium*-specific adaptation suitable for exploration as a metabolic vulnerability for parasite-directed antimalarial therapy.

## RESULTS

### Mitochondrial ACP is essential for *P. falciparum* despite loss of FASII in this organelle

The *P. falciparum* genome encodes two ACP homologs that include the well-studied protein targeted to the apicoplast (aACP, PF3D7_0208500) and a second homolog annotated as a mitochondrial ACP (mACP, PF3D7_1208300). Unlike the apicoplast ACP, which retains the conserved Ser for 4-Ppant attachment, mACP has curiously replaced this Ser with a Phe residue that cannot be modified by a 4-Ppant group (Fig. 1A and Fig. S1A). This Ser-to-Phe substitution in mACP is consistent with the loss of mitochondrial FASII and the ACP-modifying phosphopantetheine transferase enzyme (Table S1) and suggested a non-canonical function for mACP.

**Fig. 1.**
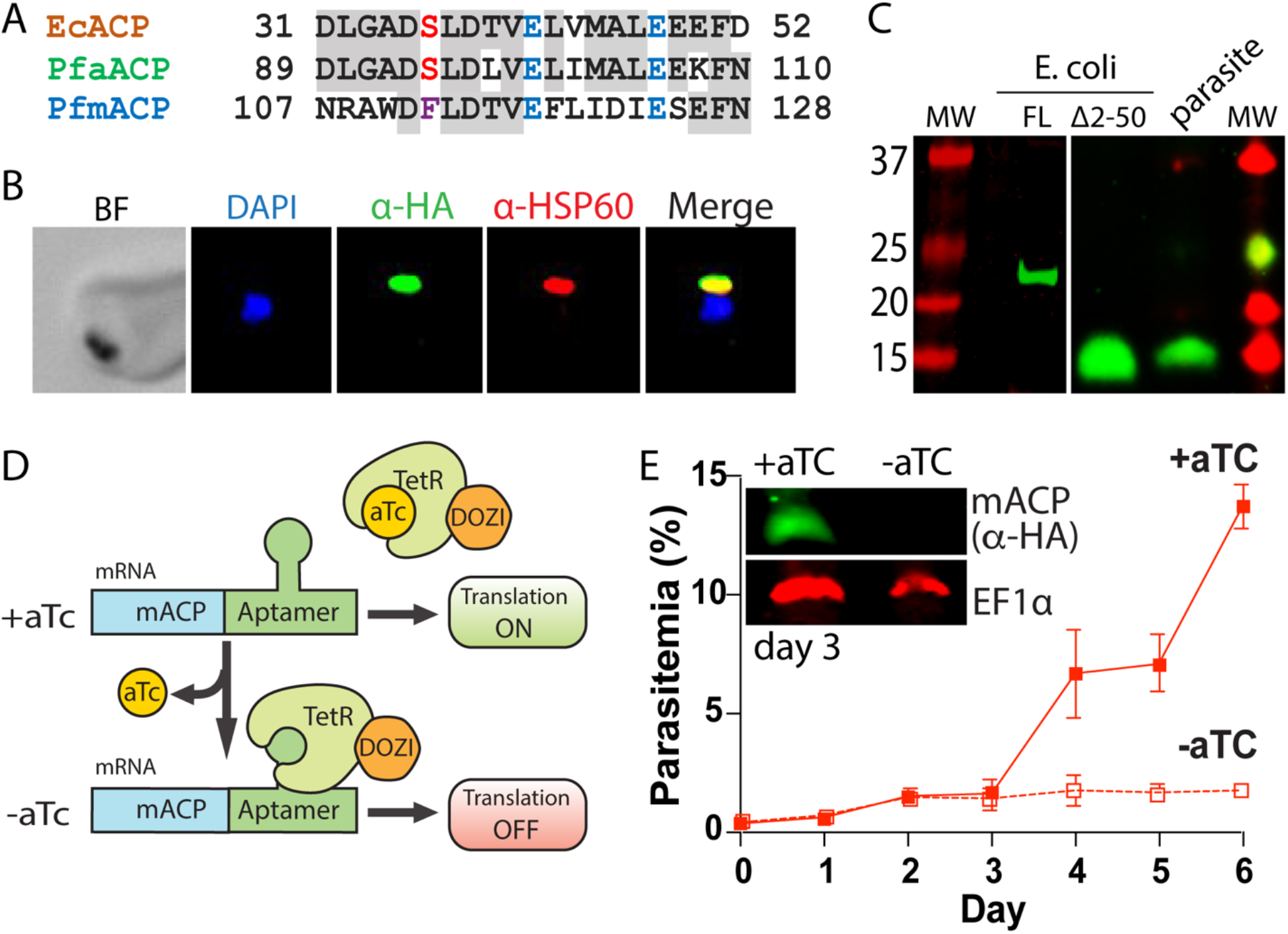
Mitochondrial ACP is essential for blood-stage *P. falciparum*. (A) Partial sequence alignment of acyl carrier protein homologs from *E. coli* and the *P. falciparum* apicoplast (aACP) and mitochondrion (mACP). The conserved Ser residue that is modified by a 4-Ppant group is in red, the divergent Phe residue of mACP is in purple, and conserved Glu residues are in blue. (B) Immunofluorescence microscopy of fixed Dd2 parasites episomally expressing mACP-HA_2_ and stained with DAPI (nucleus, blue), anti-HA (green), and anti-HSP60 (mitochondrion, red) antibodies (BF = bright field). (C) Anti-HA western blot (WB) analysis of cellular lysates of *E. coli* expressing full-length (FL) or truncated (Δ2-50) mACP-HA_2_ and Dd2 *P. falciparum* parasites episomally expressing full-length mACP-HA_2_. The left and right images are from different WB experiments (10% gels) that were aligned by molecular weight (MW) markers. (D) Schematic depiction of protein translational regulation using the aptamer/TetR-DOZI system. Normal protein expression occurs in the presence but not absence of anhydrotetracycline (aTc). (E) Continuous growth assay of synchronous mACP-HA/FLAG-aptamer/TetR-DOZI parasites in the presence or absence of aTc. Data points and error bars are the average and standard deviation from two biological replicates. Inset is an anti-HA and anti-EF1α western blot of parasite lysates harvested on day 3 of the continuous growth assay.

To test if mACP localized to the parasite mitochondrion, we created a Dd2 *P. falciparum* line that episomally expressed mACP fused to a C-terminal dual hemagglutinin (HA) tag. Immunofluorescence analysis of this mACP-HA_2_ line revealed strong co-localization between mACP-HA_2_ and the mitochondrial marker, HSP60 (Fig. 1B and Fig. S2). On the basis of this colocalization, we conclude that mACP is indeed targeted to the parasite mitochondrion, as predicted. Full-length mACP-HA_2_ has a predicted molecular weight (MW) ∼21 kDa, but SDS-PAGE and western blot (WB) analysis of immunoprecipitated mACP-HA_2_ indicated that this protein migrated with an apparent molecular weight ∼15 kDa (Fig. 1C). This lower size suggested that mACP was likely to be post-translationally processed upon mitochondrial import, which would be consistent with known processing of mitochondrial ACP in yeast and humans (28, 29).

We extended our WB analysis to include mACP-HA_2_ that was heterologously expressed in *E. coli*, where proteolytic processing is not expected. The recombinant, full-length mACP-HA_2_ expressed in bacteria migrated with an apparent MW close to 21 kDa, as expected. Based on sequence homology, the known ACP-processing sites in yeast and humans (28, 29), and mass spectrometry analysis of parasite-expressed mACP (Fig. S1B), we identified Leu-51 as the possible N-terminus of mature mACP in *P. falciparum*. We therefore cloned and bacterially expressed a truncated mACP-HA_2_ construct beginning with Leu-51. This truncated mACP protein co-migrated by SDS-PAGE with mACP-HA_2_ expressed in parasites (Fig. 1C), suggesting that Leu-51 is at or near the N-terminus of mature mACP in parasites. We conclude that mACP is proteolytically processed upon import into the *P. falciparum* mitochondrion.

To directly test if mACP is essential for blood-stage parasite growth and viability, we used CRISPR/Cas9 to tag the mACP gene in Dd2 parasites to encode a C-terminal HA-FLAG epitope tag and the aptamer/TetR-DOZI system that enables anhydrotetracycline (aTc)-dependent control of protein expression (Fig. 1D) (30). Correct integration into the mACP locus was confirmed by PCR analysis of polyclonal and clonal parasite lines (Fig. S3). To evaluate mACP knockdown and its impact on parasite growth, we synchronized parasites to the ring stage, split these parasites into two equal populations ±aTc, and monitored parasite growth over multiple 48-hour growth cycles. Parasites grew indistinguishably ±aTc for the first 3 days. However, parasites grown without aTc displayed a major growth defect on day 4 in the third intraerythrocytic lifecycle (Fig. 1E), similar to prior studies with this knockdown system (30). Blood-smear analysis on day 5 indicated wide-spread parasite death (Fig. S4). Western blot analysis of parasite samples harvested in the second cycle, after three days of growth -aTc, indicated robust knockdown of mACP expression relative to parasites grown +aTc (Fig. 1E). We conclude that mACP is essential for *P. falciparum* parasites during blood-stage growth.

### Mitochondrial ACP binds the Isd11-Nfs1 complex required for Fe-S cluster biogenesis

Recent studies in yeast and human cells have revealed that mitochondrial ACP binds to a variety of small, 3-helical adapter proteins that bear a conserved Leu-Tyr-Arg (LYR) sequence motif on their N-terminal helix (24, 31). The Leu mediates an intramolecular helical-helical contact while the Tyr and Arg side-chains interact with conserved acidic residues on ACP (25, 27, 32). These LYR-motif proteins mediate diverse mitochondrial processes that include respiratory chain assembly, Fe-S cluster biogenesis, and ribosomal translation (20, 21, 23, 26). Given the strong conservation of mitochondrial LYR proteins and ACP interactions across eukaryotes, we reasoned that the essential function of *P. falciparum* mACP might involve interaction with a conserved LYR protein.

Using the sequences of known LYR proteins from yeast and humans as bait (31), we conducted a BLAST search of the *P. falciparum* genome to look for homologs. We identified Isd11 (PF3D7_1311000) as the only LYR-protein homolog retained by parasites (Table S2). *P. falciparum* Isd11 is 29% identical to human Isd11 (also known as LYRM4), including conservation of the LYR sequence motif near the N-terminus (Fig. 2A). The parasite mACP retains the conserved Glu residues (5 and 11 amino acids C-terminal to the modified Ser in canonical ACPs) known to interact with the Tyr and Arg residues of the LYR motif on Isd11 (Fig. 1A) (25, 27). In yeast and humans, Isd11 directly binds and stabilizes Nfs1, a mitochondrial cysteine desulfurase required for biogenesis of Fe-S clusters (33). Mitochondrial ACP in these organisms directly binds Isd11 to stabilize the mACP-Isd11-Nfs1 complex (Fig. 2B), with loss of ACP and/or Isd11 resulting in Nfs1 instability and defective Fe-S cluster biogenesis (20, 34-36). *P. falciparum* retains a mitochondrial Nfs1 homolog (PF3D7_0727200) that is predicted from genome-wide knock-out studies to be essential (19, 37) but whose role in Fe-S cluster biogenesis has not been studied. Collectively, conservation of Isd11 and Nfs1 in *P. falciparum*, the conserved role for ACP in stabilizing the mACP-Isd11-Nfs1 complex in other eukaryotes, and retention of the LYR sequence motif in *P. falciparum* Isd11 strongly suggested that mACP was likely to interact with Isd11 to form the mACP-Isd11-Nfs1 complex in parasites.

**Fig. 2.**
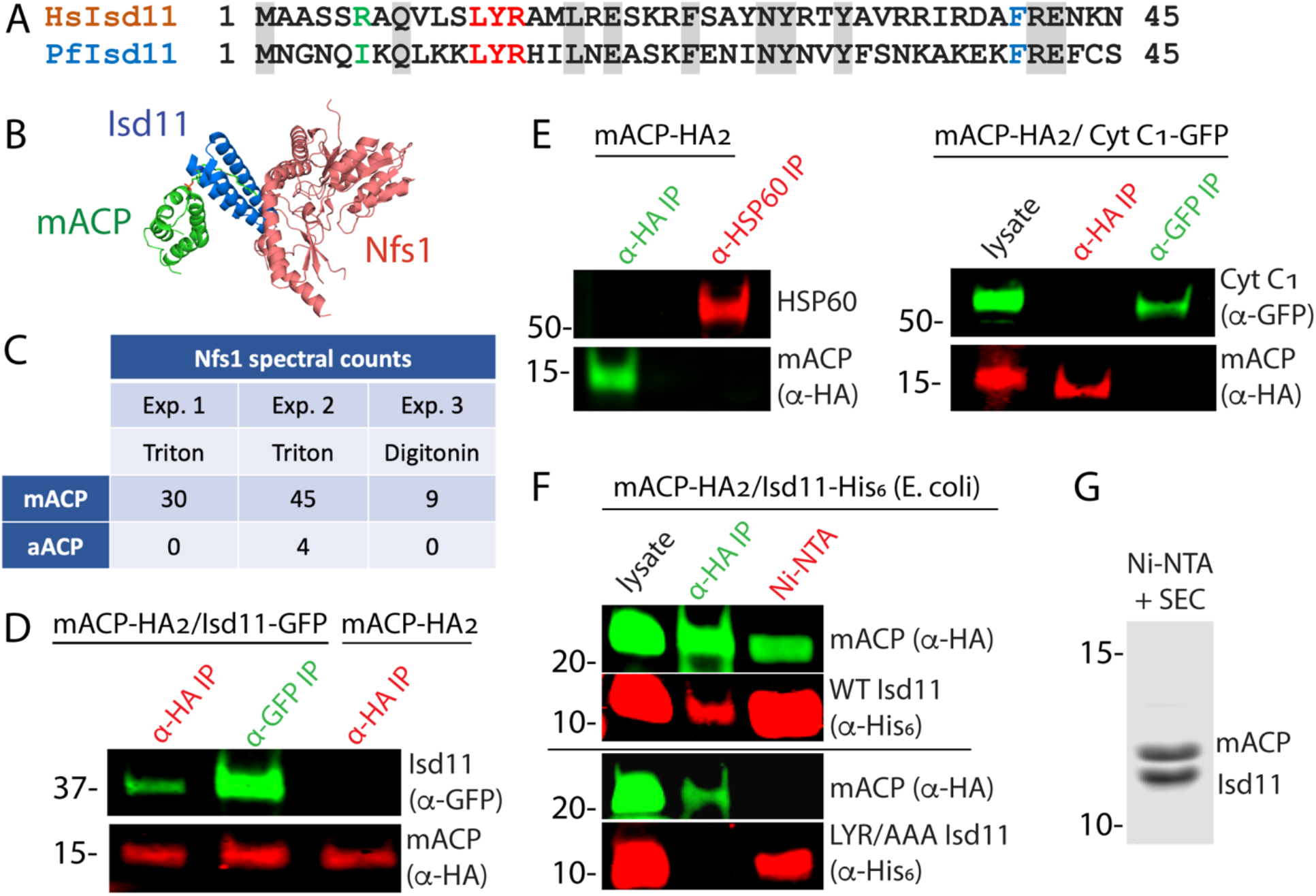
Mitochondrial ACP binds the Isd11-Nfs1 complex. (A) Partial sequence alignment of human and *P. falciparum* Isd11. The conserved LYR sequence motif is in red, the conserved Phe residue is in blue, and residue 6 is in green. (B) X-ray structural model of the mACP-Isd11-Nfs1 complex (PDB entry 5USR). For simplicity, only a single copy of each protein is shown, rather than the functional dimer (27). (C) Table of spectral counts for *P. falciparum* Nfs1 detected by tandem mass spectrometry for anti-HA immunoprecipitation studies of lysates from Dd2 parasites episomally expressing mACP-HA_2_ or aACP-HA_2_. Parasites were lysed in either Triton X-100 or digitonin. Similar spectral counts for mACP and aACP bait proteins were detected in each experiment (Table S3). (D) Anti-HA or anti-GFP co-immunoprecipitation/western blot studies of Dd2 parasites episomally expressing mACP-HA_2_ with or without Isd11-GFP. (E) Anti-HA, anti-HSP60, or anti-GFP co-IP/WB studies of Dd2 parasites episomally expressing mACP-HA_2_ with or without cyt c_1_-GFP. (F) Anti-HA co-IP or nickel-nitrilotriacetic acid (Ni-NTA) pulldown and WB studies of *E. coli* bacteria recombinantly expressing full-length mACP-HA_2_ and Isd11-His_6_ (WT or LYR/AAA mutant) and probed with anti-HA and anti-His_6_ antibodies. All lanes are from the same blot and were processed identically but were cropped from non-contiguous lanes. (G) Coomassie-stained SDS-PAGE gel of recombinant His_6_-Isd11 and Δ2-50 mACP-HA_2_ purified from *E. coli* by Ni-NTA pulldown of His_6_-Isd11 and size-exclusion chromatography (SEC) after removal of affinity tags. Full images for all WB experiments are shown in Fig. S6.

To identify protein interaction partners of mACP, we used anti-HA immunoprecipitation (IP) to isolate mACP-HA_2_ from parasites followed by tandem mass spectrometry (MS) to identify parasite proteins that co-purified with mACP. In multiple independent pull-downs, we identified Nfs1 as strongly enriched in the mACP-HA_2_ sample compared to a negative control sample (anti-HA IP of aACP-HA_2_) (Fig. 2C and Fig. S5). We did not observe peptides corresponding to Isd11 in mACP-HA_2_ IP/MS samples. However, its absence is not unexpected given the short Isd11 length (87 residues) that leads to comparatively few tryptic peptides.

To directly test for mACP interaction with Isd11, we transfected the mACP-HA_2_/Dd2 line with an episome encoding expression of Isd11 with a C-terminal GFP tag that was previously used to localize Isd11 to the mitochondrion (37). Using this Dd2 line expressing mACP-HA_2_ and Isd11-GFP, we performed reciprocal IP/WB experiments that confirmed stable interaction of these two proteins in parasites (Fig. 2D and Fig. S6A). This interaction appeared to be specific, as mACP-HA_2_ did not co-IP with the abundant mitochondrial chaperone, HSP60, or with GFP-tagged cytochrome c_1_ (Fig. 2E and Fig. S6B & S6C). Because these IP experiments cannot distinguish whether mACP binding to Isd11 is direct or mediated by other parasite proteins, we turned to heterologous studies in *E. coli*. Reciprocal IP experiments with lysates from bacteria co-expressing *P. falciparum* mACP-HA_2_ and Isd11-His_6_ confirmed stable interaction of these proteins in the absence of other parasite-specific factors (Fig. 2F). In experiments with Isd11 LYR mutants, however, this interaction was strongly reduced (YR/AA) or eliminated (LYR/AAA), despite robust expression of all proteins (Fig. 2F and Fig. S6D). Clarified lysates from bacteria co-expressing truncated (Δ2-50) mACP-HA_2_ and Isd11-His_6_ were also fractionated by passage over Ni-NTA resin to selectively pull down His-tagged Isd11 and interactors followed by size-exclusion chromatography to separate individual complexes. Analysis by Coomassie-stained SDS-PAGE and tandem mass spectrometry indicated robust isolation of a 1:1 complex containing Isd11 and mACP (Fig. 2G and Fig. S7). We conclude that mACP directly binds to Isd11 and that this interaction requires the LYR motif.

Yeast and human Isd11 retain a conserved Arg six residues upstream of the LYR motif (Fig. 2A) that electrostatically stabilizes the negatively charged oxygen atoms of the Ppant group in the ACP-Isd11 complex (25, 27, 32), and mutation of this Arg ablates binding of human Isd11 to ACP (24). Because parasite mACP lacks the negatively charged Ppant group and has replaced it with a hydrophobic Phe, retention of Arg-6 by *P. falciparum* Isd11 would be predicted to destabilize binding of these two proteins. The parasite Isd11, however, has replaced Arg-6 with an Ile residue (Fig. 2A) expected to more favorably interact with the hydrophobic surface of mACP created by Ser-to-Phe substitution. These sequence features suggest that parasite Isd11 has co-evolved with mACP to optimize formation of the mACP-Isd11 complex in *P. falciparum* in the absence of FASII and an acyl-Ppant group on mACP. Consistent with this model, parasite Isd11 did not appear to interact with *E. coli* ACP, which retains the acyl-Ppant modification, in contrast to human Isd11 that robustly binds bacterial ACP when expressed in *E. coli* (25, 27, 36).

### Structural modeling of the mACP-Isd11 interface

Association of Isd11 and mACP in yeast and human cells depends critically on the acyl-Ppant group of mACP (Fig. 3A), with little or no association observed in these cells between Isd11 and the mACP Ser-to-Ala mutant that lacks a Ppant group (20, 21, 24). As an initial step to understand the divergent molecular features of the *P. falciparum* mACP-Isd11 interface that stabilize complexation in the absence of an acyl-Ppant group, we generated a homology model of this complex using published structures and then carried out energy optimization and refinement using the Rosetta software (38). The resulting low-energy structural model suggested that the unusual Phe113 residue of mACP partitions into a hydrophobic pocket at the mACP-Isd11 interface composed of the aliphatic side-chain methylene groups of Lys10 and Arg14 and the phenyl ring of Tyr13 of Isd11 as well as Val117 on mACP (Fig. 3B).

**Fig. 3.**
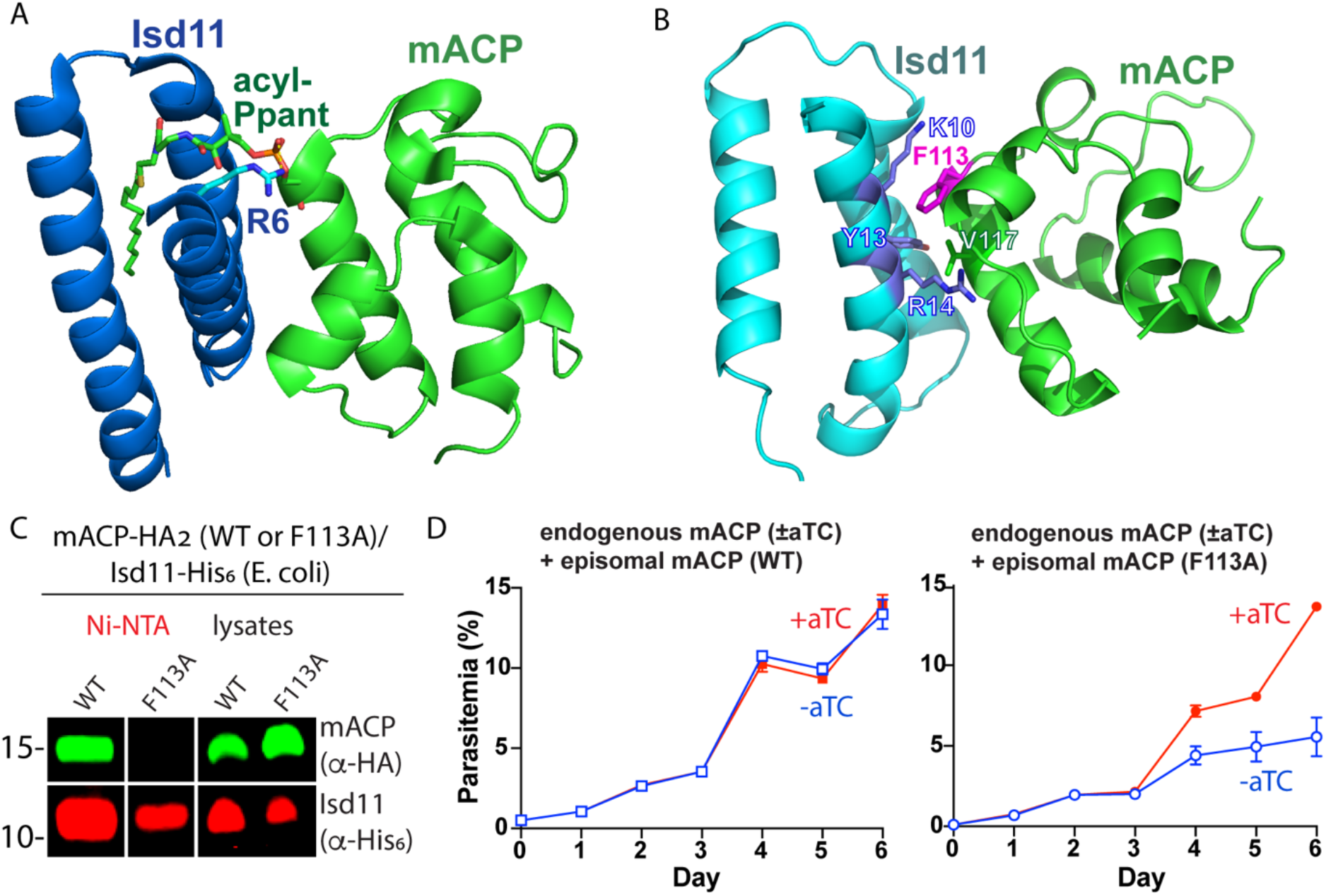
Structural modeling and functional tests of the divergent Phe113 residue of parasite mACP. (A) X-ray crystallographic structure of human Isd11 bound to *E. coli* ACP highlighting the central role of the acyl-Ppant group of ACP and R6 residue of Isd11 in stabilizing this complex (PDB entry 5USR). (B) Rosetta-based, energy-minimized structural model of the *P. falciparum* mACP-Isd11 interface. (C) Nickel-nitrilotriacetic acid (Ni-NTA) pulldown and WB studies of *E. coli* bacteria recombinantly expressing Δ2-50 mACP-HA_2_ (WT or F113A mutant) and Isd11-His_6_ and probed with anti-HA and anti-His_6_ antibodies. All lanes are from the same blot and were processed identically but were cropped from non-contiguous lanes. The uncropped WB is shown in Fig. S6E. (D) Continuous growth assays of synchronous mACP-aptamer/TetR-DOZI parasites episomally expressing WT or F113A mACP-HA_2_ and grown in the presence or absence of aTc. Data points and error bars are the average and standard deviation from two biological replicates. The presence of an HA epitope on both the endogenous and episomal mACP prevented determination of endogenous mACP expression differences ±aTc by WB.

To test if the divergent Phe113 residue of mACP contributes substantially to its association with Isd11, we heterologously co-expressed Isd11-His_6_ and Δ2-50 mACP-HA_2_ (WT or the F113A mutant) in *E. coli* and observed robust expression of all proteins in bacterial lysates. We then used Ni-NTA to selectively pull down His-tagged Isd11 and observed that WT mACP-HA_2_ but not the F113A mutant co-purified with Isd11 (Fig. 3C and S6E). These results indicate that mutation of Phe113 to Ala substantially weakens the mACP-Isd11 interaction, supporting the conclusion that Phe113 contributes to the mACP-Isd11 binding interface in the mACP-Isd11-Nfs1 complex as suggested by structural modeling (Fig. 3B).

To test the functional impact of the Phe113Ala mACP mutation on parasite growth, we transfected our mACP knockdown parasite line with episomes encoding either WT or Phe113Ala mACP. Parasites expressing a second-copy of WT mACP showed identical growth in the presence or absence of aTc. In contrast, parasites that episomally expressed the Phe113Ala mACP mutant showed strongly diminished growth upon aTc removal and loss of endogenous mACP (Fig. 3D). These results support the conclusion that the Phe residue of mACP plays a key role in stabilizing the mACP-Isd11 association required for Nfs1 stability and function.

### Loss of mACP destabilizes Nfs1 and the Rieske protein in ETC complex III

Nfs1 stability in other eukaryotes depends critically on its association with Isd11 and ACP, with loss of ACP resulting in degradation of Nfs1 and lethal defects in Fe-S cluster biogenesis (20, 25, 27, 34, 36). Loss of Nfs1 upon mACP knockdown would therefore be sufficient to explain mACP essentiality in *P. falciparum*. To test the impact of mACP knockdown on Nfs1 protein levels in parasites, we used a commercial anti-Nfs1 antibody (Abcam 229829) raised against a 250-amino acid region of human Nfs1 that is 55% identical to *P. falciparum* Nfs1 and which selectively recognized recombinant *P. falciparum* Nfs1 expressed in *E. coli* (Fig. S6F). We grew our mACP aptamer/TetR-DOZI parasites ±aTc, harvested them at the end of the second intraerythrocytic growth cycle before parasite death, and analyzed expression of Nfs1 by western blot using this Nfs1 antibody.

In parasites grown +aTc, the anti-Nfs1 antibody recognized a band at ∼55 kDa (Fig. 4A) that is slightly lower than the ∼63 kDa band recognized by this antibody for full-length Nfs1 expressed in *E. coli* (Fig. S6C). This observation suggests that Nfs1 is processed upon mitochondrial import in *P. falciparum*, which is consistent with observed processing of Nfs1 in yeast (28). Western blot analysis of parasites grown -aTc failed to detect this 55 kDa band, strongly suggesting that Nfs1 stability is tightly coupled to mACP expression and that loss of mACP results in Nfs1 degradation (Fig. 4A). Loss of Nfs1 upon mACP knockdown suggests that parasite death is due to dysfunction in one or more Fe-S cluster-dependent pathways. We set out to understand which pathway defect(s) might explain parasite death.

**Fig. 4.**
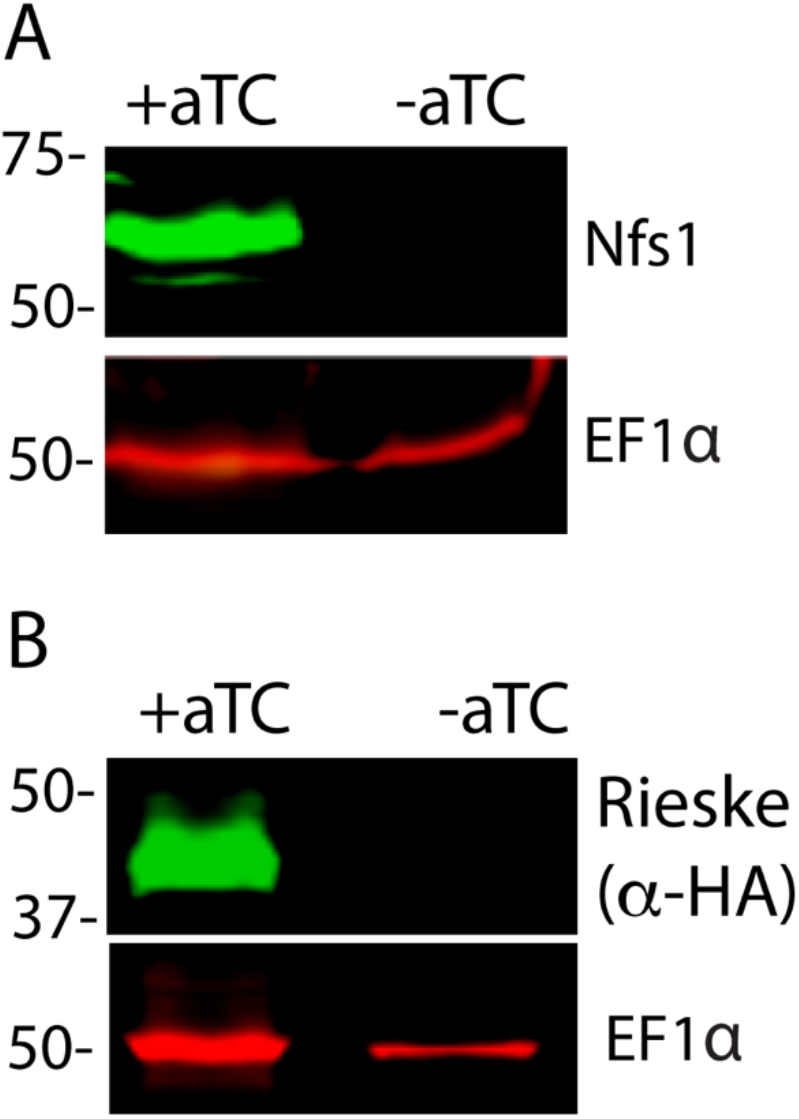
Loss of mACP destabilizes Nfs1 and the Rieske protein. Western blot analysis of lysates from mACP-aptamer/TetR-DOZI parasites episomally expressing Rieske-HA_2_ that were grown ±aTc for 3 days and probed for EF1α and (A) Nfs1 or (B) HA.

The mitochondrial iron-sulfur cluster (ISC) biogenesis pathway supplies Fe-S clusters to various mitochondrial proteins and has been suggested from studies in yeast to produce a key intermediate that is exported out of the organelle and elaborated by cytoplasmic proteins to assemble and deliver Fe-S clusters to client proteins in the cytoplasm and nucleus (34, 39-41). Multiple Fe-S cluster proteins function within the *P. falciparum* mitochondrion (42). Aconitase (PF3D7_1342100) and succinate dehydrogenase (PF3D7_1212800) are dispensable for blood-stage parasite growth (8). Ferredoxin (PF3D7_1214600) is expected to be essential (19), but its main role is to provide electrons for the ISC pathway (43). Class I fumarate hydratase (PF3D7_0927300), which functions in the TCA cycle but may also contribute to purine scavenging, was refractory to disruption in *P. falciparum* (8) but successfully deleted in *P. berghei* in a mouse strain-dependent manner (44). The Rieske protein (PF3D7_1439400), which is an essential component of ETC complex III (45), is the only known mitochondrial Fe-S cluster protein that has an unequivocally essential function apart from the ISC pathway. We therefore focused on understanding the effect of mACP knockdown on the Rieske protein, as a prior study in yeast showed that mACP knockdown resulted in loss of Rieske (20).

To probe the status of Rieske upon mACP knockdown in parasites, we transfected the mACP-aptamer/TetR-DOZI parasites with an episome encoding Rieske-HA_2_. A band corresponding to the epitope-tagged Rieske was detected ∼43 kDa in western blot analysis of parasites grown +aTc. This band, however, was undetectable in parasites grown 72 hours in -aTc conditions to downregulate mACP expression (Fig. 4B). These results indicate that knockdown of mACP and subsequent degradation of Nfs1 result in loss of the Fe-S cluster-dependent Rieske protein.

### Loss of mACP causes ETC failure and sensitizes parasites to mitochondrial depolarization by proguanil

Since the Rieske protein is an essential electron-transfer component of ETC complex III (45), we reasoned that loss of mACP and Rieske likely results in ETC failure. In blood-stage parasites, which rely on glycolysis rather than oxidative phosphorylation for ATP synthesis, the essential function of the mitochondrial ETC is to oxidatively recycle the ubiquinone cofactor used by several dehydrogenases, of which dihydroorotate dehydrogenase is most critical (1, 46). Prior work has shown that exogenous addition of the soluble ubiquinone analog, decyl-ubiquinone (dQ), rescues parasites from ETC dysfunction caused by the ETC complex III inhibitor, atovaquone (46). We posited that it might be possible to rescue parasites with dQ from mACP knockdown if the dominant cause of parasite death were ETC failure.

As a positive control and to provide a basis for comparison, we first confirmed that 15 µM exogenous dQ substantially rescued parasites over multiple intraerythrocytic cycles from growth inhibition by 100 nM atovaquone (Fig. 5A), as previously reported (46). To test dQ rescue of parasites upon mACP knockdown, we synchronized the mACP-aptamer/TetR-DOZI parasites and monitored their growth ±aTc and ±dQ for several intraerythrocytic cycles. As before, -aTc parasites grew normally for three days but failed to expand into the third growth cycle on day four (Fig. 5A). Addition of dQ rescued parasite growth in the third growth cycle in -aTc conditions, similar to dQ rescue of parasites from atovaquone. However, in contrast to indefinite parasite rescue from atovaquone, dQ only rescued the third-cycle growth of parasites cultured -aTc, which failed to expand further and began to die off on day six in the fourth intraerythrocytic cycle (Fig. 5A). On the basis of dQ rescue, we conclude that ETC failure is the immediate cause of parasite death upon mACP knockdown. However, the inability of dQ to rescue parasites beyond the third cycle indicates that additional dysfunctions beyond the mitochondrial ETC contribute to parasite death on a longer timescale. These additional defects may involve mitochondrial fumarate hydratase and/or essential Fe-S cluster proteins in the cytoplasm and nucleus that depend on Nfs1 function (see Discussion below).

**Fig. 5.**
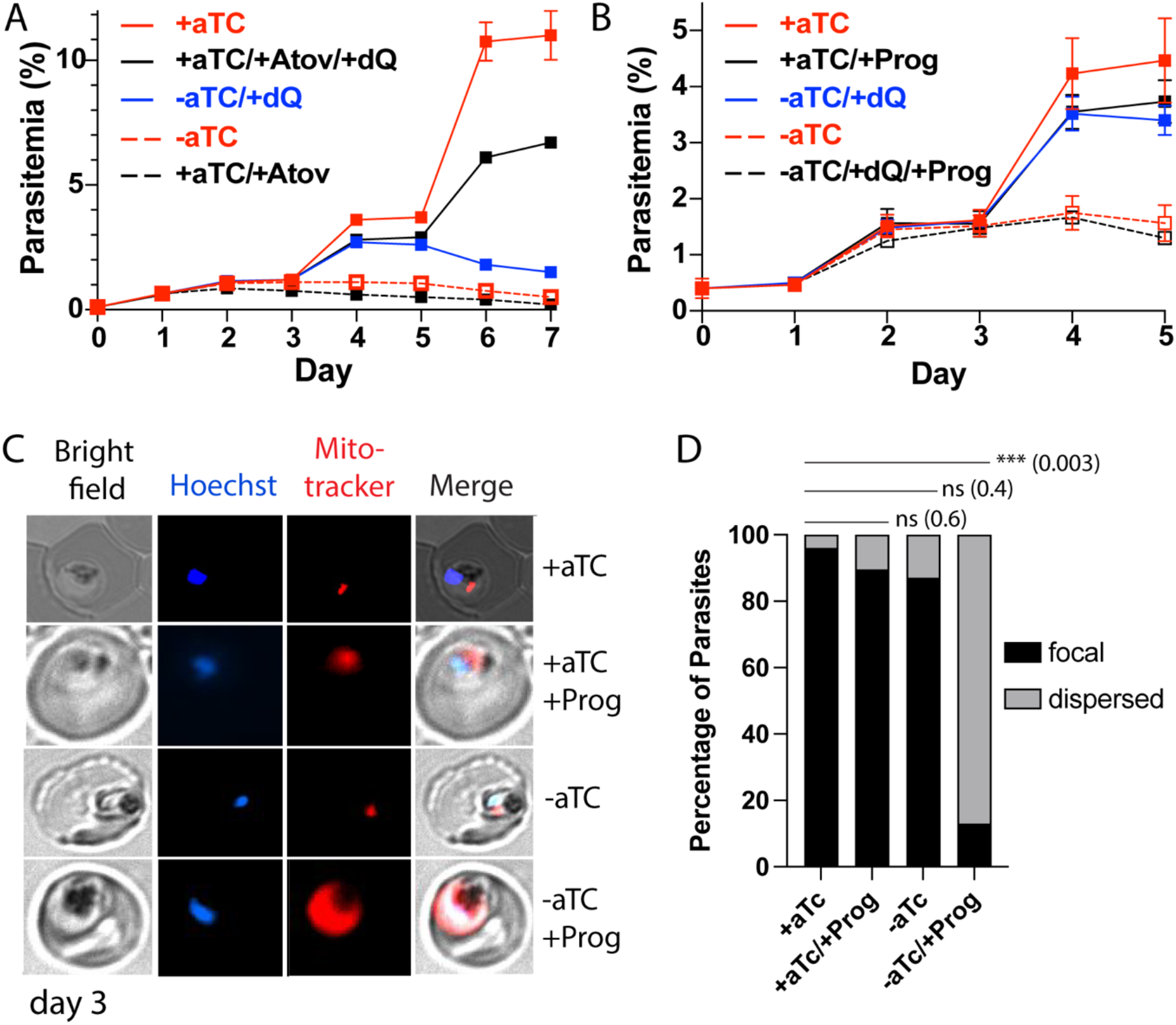
Loss of mACP causes ETC failure and sensitizes parasites to proguanil. Continuous growth assays of synchronous mACP-aptamer/TetR-DOZI parasites grown (A) ±aTc, ±atovaquone (Atov, 100 nM), ±decyl-ubiquinone (dQ, 15 µM), and (B) ±proguanil (Prog, 1 µM). Data points and error bars are the average and standard deviation from 2-3 biological replicates. (C) Fluorescence microscopy images of live mACP-aptamer/TetR-DOZI Dd2 parasites cultured 3 days ±aTc, 2 days ±5 µM proguanil, and stained with Hoechst or Mitotracker Red (10 nM). (D) Statistical analysis of MitoTracker signal for 40-50 total parasites from each condition in panel C from two independent experiments. For clarity, error bars are not shown but standard errors of the mean were ≤10% in all cases. Cell percentage differences were analyzed by two-tailed unpaired t-test (p values in parentheses, ns = not significant).

To further unravel the effect of mACP knockdown on ETC function, we next tested if loss of mACP sensitized parasites to mitochondrial depolarization by proguanil. The ETC couples ubiquinone recycling to proton translocation from the mitochondrial matrix to the inner membrane space to establish a transmembrane electrochemical potential. Prior work has shown that parasites retain a second proton-pumping mechanism that also contributes to transmembrane potential and which is inhibited by proguanil (1, 47). Inhibition of ETC function kills parasites due to defective ubiquinone recycling but does not substantially depolarize mitochondria due to activity by this second, proguanil-sensitive pathway. However, ETC dysfunction plus proguanil treatment blocks both pathways for proton pumping and causes mitochondrial depolarization, which can be visualized by a failure to concentrate charged dyes like Mitotracker within the mitochondrion that leads to dispersed dye accumulation in the cytoplasm (1, 48).

If mACP knockdown and subsequent loss of Rieske caused general ETC failure, we predicted that parasite treatment with sub-lethal proguanil would negate the ability of dQ to rescue parasite growth from mACP knockdown in the third cycle and would cause mitochondrial depolarization. We repeated the prior growth assay ±aTc and ±dQ but also included 1 µM proguanil. This proguanil concentration alone had no effect on parasite growth, as previously reported (1). However, when proguanil was combined with growth -aTc, dQ was unable to rescue parasite growth in the third cycle (Fig. 5B). Microscopy analysis of parasites on day three of the growth assay revealed that proguanil treatment selectively prevented mitochondrial accumulation of Mitotracker Red in parasites grown -aTc, strongly suggesting mitochondrial depolarization (Fig. 5C and 5D, and Fig. S8). This observation strongly supports the model that impaired ubiquinone recycling due to Rieske loss and ETC dysfunction is the immediate cause of parasite death upon mACP knockdown.

## DISCUSSION

The *P. falciparum* mitochondrion is a major antimalarial drug target, but nearly all organelle-specific inhibitors target cytochrome b in ETC complex III or dihydroorotate dehydrogenase (DHOD) that depends on complex III function (49, 50). Multiple metabolic pathways operate within the mitochondrion (3), but many are dispensable for blood-stage parasites and few essential functions beyond DHOD and ETC complexes III and IV have been identified. Iron-sulfur cluster biosynthesis is an ancient, essential mitochondrial function that has been well studied in yeast and mammalian cells but is sparsely studied in parasites (37). We have identified a divergent and essential nexus between Fe-S cluster biogenesis and an evolutionary vestige of type-II fatty acid synthesis in the *P. falciparum* mitochondrion.

### New molecular paradigm for essential ACP function without an acyl-Ppant group

Most eukaryotic cells, including fungi, plants, and animals, retain a mitochondrial FASII pathway in which the acyl-ACP intermediate has been shown to critically mediate respiratory chain assembly, Fe-S cluster biogenesis, and ribosomal translation by binding to LYR-protein assembly factors (24, 31, 51). These interactions and their dependence on ACP acylation have been proposed to constitute a regulatory feedback mechanism that couples the availability of acetyl-CoA for FASII activity to respiratory chain assembly for oxidative phosphorylation and ATP synthesis (21). Biochemical coupling of these pathways, however, has not been retained in *Plasmodium*, which has lost has lost mitochondrial FASII enzymes but retains a divergent ACP homolog incapable of 4-Ppant modification or acylation. Nevertheless, we have shown that mACP remains essential to parasites for Fe-S cluster biogenesis by binding Isd11 and stabilizing the mACP-Isd11-Nfs1 cysteine desulfurase complex via a novel interface. *P. falciparum* thus provides a new molecular paradigm for essential ACP function without an acyl-Ppant group. This discovery emphasizes the ancient, fundamental role of ACP in mitochondrial Fe-S cluster biogenesis and suggests an evolutionary driving force to retain mACP interaction with the Isd11-Nfs1 complex independent ACP’s scaffolding role in fatty acid synthesis.

Prior in vitro work has provided evidence that activity of purified Isd11-Nfs1 does not strictly require association with ACP (27, 34). However, Nfs1 is unstable in its native mitochondrial context in the absence of ACP, as shown herein for *P. falciparum* and in prior studies of yeast (20, 52). Coupling Nfs1 stability (via Isd11) to ACP acylation in eukaryotes that retain mitochondrial FASII has been proposed as a mechanism to up-and down-regulate Fe-S cluster biogenesis congruent with nutrient availability and cellular Fe-S cluster needs for respiration and growth (34). This mechanism, however, cannot explain mitochondrial retention of an acylation-incompetent ACP homolog in *Plasmodium*. ACP may therefore play additional functional and/or regulatory roles that are essential for mitochondrial Fe-S cluster biogenesis and utilization in cells, perhaps involving metal-ion binding and sensing by ACP (e.g., iron or zinc) (32, 53) or interactions with other mitochondrial networks akin to broad ACP interactions observed in *E. coli* (54). In this regard it is interesting to note that anaerobic eukaryotic parasites like *Giardia* and *Cryptosporidium* lack a mitochondrion but retain a primitive mitosome for Fe-S cluster biogenesis that lacks ACP and Isd11 homologs in this compartment, where Nfs1 appears to function autonomously (55, 56).

### Implications for parasites

Loss of mitochondrial FASII but retention of a Ppant-independent mACP lacking the conserved Ser appears to be widespread among Apicomplexan parasites, including *Toxoplasma* and *Babesia*. Outside the phylum Apicomplexa, the only genomic evidence we found for a divergent ACP lacking the conserved Ser was in the related photosynthetic chromerid, *Vitrella brassicaformis*, but not in its algal cousin, *Chromera velia*. Thus, the adaptation of mitochondrial ACP to function without a Ppant group or acylation likely occurred on a similar evolutionary time-frame as the loss of plastid photosynthesis that accompanied the appearance of Apicomplexan parasitism (57).

Our study indicates that mitochondrial ACP function without an acyl-Ppant group is a parasite-specific adaptation that differentiates Apicomplexan parasites from humans. Nevertheless, like yeast and human cells (20, 24), we observed that loss of ACP destabilizes Nfs1 to block Fe-S cluster assembly, impairs the stability and function of the key Rieske subunit of ETC complex III, and leads to parasite death (Fig. 6). In yeast and mammalian cells, mACP plays a second role in Rieske maturation by binding to Mzm1, an LYR-protein chaperone that stabilizes Rieske upon import into the mitochondrial matrix to receive its Fe-S cluster for subsequent insertion into ETC complex III by the AAA-ATPase, BCS1 (24, 58). *P. falciparum*, however, appears to lack an Mzm1 homolog (Table S2), suggesting that parasites have evolved compensatory mechanisms for Rieske maturation in the absence of this ACP-dependent chaperone.

**Fig. 6.**
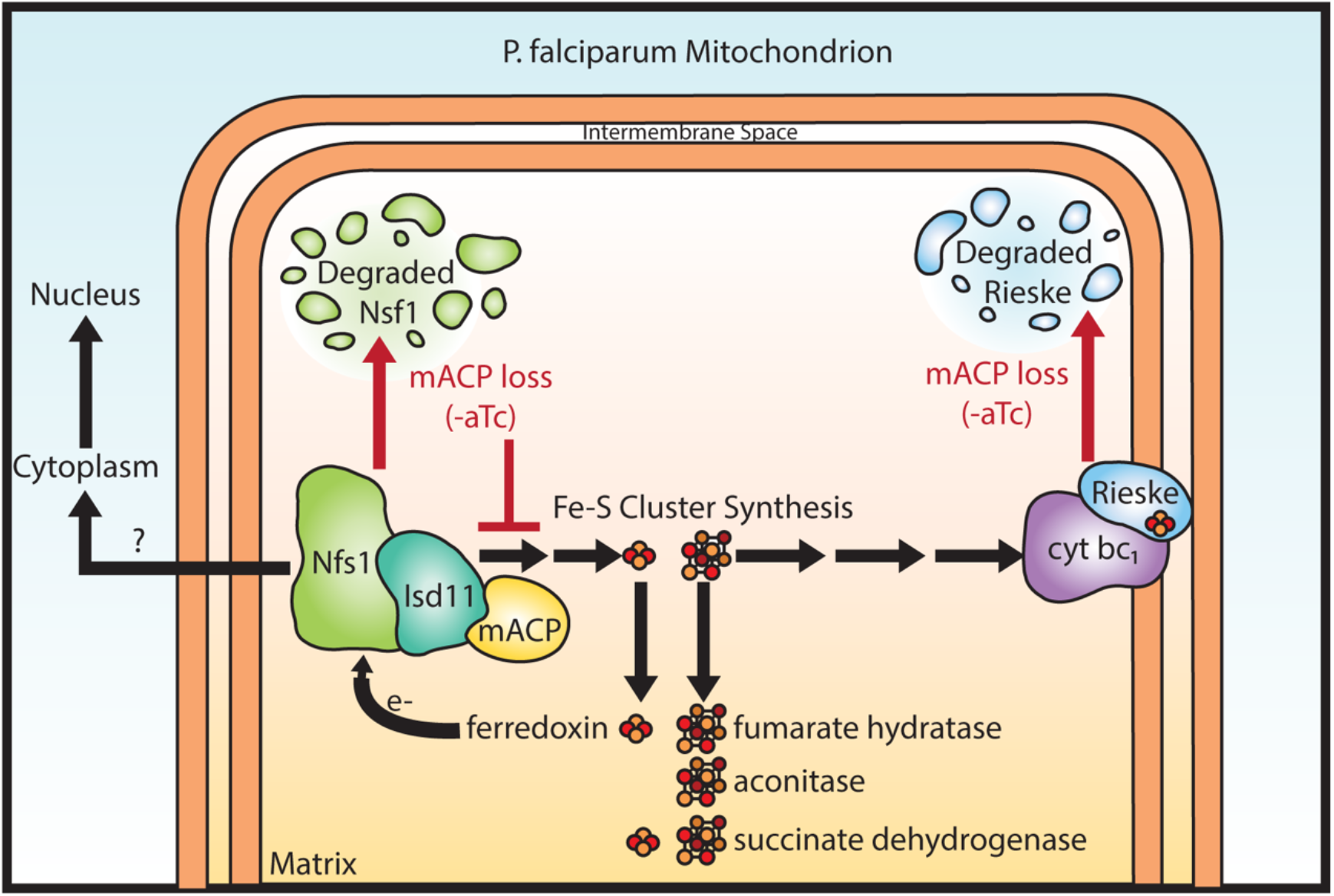
Schematic model for the impact of mACP knockdown (-aTc) on Fe-S cluster metabolism in *P. falciparum* parasites. The question mark indicates uncertainty about in the functional role of mitochondrial Nfs1 in supporting cytoplasmic Fe-S cluster biogenesis in parasites. For simplicity, the mACP/Isd11/Nfs1 complex is shown with only one monomer for each protein, rather than the functional dimer (27). Illustration by Megan Okada.

Exogenous dQ rescued parasite growth upon mACP knockdown for one intraerythrocytic cycle but did not rescue growth on a longer time-scale. This short-term rescue indicates that respiratory failure is the immediate cause of *P. falciparum* death upon loss of mACP (1, 46). However, defects in essential processes beyond the mitochondrial ETC contribute to parasite death on longer timescales. These defects may involve mitochondrial fumarate hydratase (8) and/or essential Fe-S cluster proteins in the cytosol and nucleus that depend on Nfs1 activity (Fig. 6), including Rli1 required for cytoplasmic ribosome assembly and Rad3 and Pri2 that are essential for nuclear DNA metabolism (59). In yeast, the mitochondrial transporter Atm1 has been proposed to export a key sulfur-containing intermediate required for cytosolic Fe-S cluster assembly (39, 56). In *P. falciparum*, the Mdr2 transporter (PF3D7_1447900) is 34% identical to yeast Atm1 and has been proposed to perform an analogous function in cytosolic Fe-S cluster assembly (3). However, Mdr2 appears to be dispensable for blood-stage parasites (60), and its possible mitochondrial-export role in cytoplasmic Fe-S cluster biogenesis remains untested. We also note prior studies in mammalian cells which suggest that cytoplasmic Fe-S cluster synthesis in these cells may be independent of mitochondrial Nfs1 activity (61, 62). Cytoplasmic Fe-S cluster metabolism in *Plasmodium* is sparsely studied, and its functional dependence on mitochondrial ISC proteins remains a key frontier to further test and understand.

The mitochondrial ETC is essential for malaria parasite viability in all life-cycle stages (63). The unique molecular features of *P. falciparum* mACP and its interaction with Isd11 that underpin essential functions of the ETC and broader Fe-S cluster utilization suggest the possibility of targeting this complex for antimalarial therapy. We have on-going structural and biochemical studies to assess if parasite Isd11 retains a vestigial acyl-pantetheine binding pocket and to test if parasite-specific features of the mACP-Isd11 interaction can be selectively disrupted via small-molecule inhibitors that mimic the acyl-pantetheine group and/or broader hydrophobic features of this protein-protein interface that are distinct from the human complex. On the basis of molecular conservation in other Apicomplexan parasites, such inhibitors would likely function against other human pathogens such as *Toxoplasma* and *Babesia*.

## MATERIALS AND METHODS

Key experimental materials and methods are described below. In the SI Appendix, additional details are provided in SI Methods for cloning of parasite genes into expression and genome-editing vectors, sample processing steps and instrument settings for mass spectrometry experiments, and validation of the anti-Nfs1 antibody. We also provide a list of cloning primers used and the accession codes for the known LYR protein homologs from yeast and humans. Vendor and product information is also listed for key materials when mentioned for the first time. Descriptions of statistical analyses and samples sizes are indicated below, in figure legends, and in SI Methods.

### Parasite growth assays

All parasite studies were performed with *P. falciparum* Dd2 parasites cultured in RPMI-1640 medium as previously described (64). CRISPR/Cas9 was used to tag the mACP gene in Dd2 parasites to encode a C-terminal HA-FLAG epitope tag and the aptamer/TetR-DOZI system (30), whose integration was validated by PCR. Growth assays were performed ±aTc by synchronizing parasite by 5% D-sorbitol, monitoring culture parasitemia by acridine orange staining and flow cytometry of ≥2 biological replicate samples, and reporting the parasitemia average and standard deviation. Decyl-ubiquinone was used in chemical rescue experiments at 15 µM final concentration and was added ±1 µM proguanil at the beginning of growth assays.

### Western blot analyses

Parasite sample were harvested by 0.05% saponin treatment. For whole-parasite experiments, saponin pellets were dissolved in 60 µl SDS sample buffer containing beta-mercaptoethanol. For immunoprecipitation experiments, parasite pellets from 50-mL cultures were lysed in 1 mL 1% triton or digitonin and clarified by centrifugation. Lysate supernatants were incubated with 30 µl of pre-equilibrated Pierce anti-HA magnetic beads, washed 3X with Tris-buffered saline + 0.05% Tween-20, eluted with 8M urea, and diluted with 5X SDS sample buffer. For bacteria expression studies, BL21/DE3 *E. coli* were transformed with pET plasmids encoding cloned *P. falciparum* mACP-HA_2_ (full-length or the Δ2-50 truncation, WT or F113A mutant), Isd11-His_6_ (WT, YR/AA, or LYR/AAA mutants), or full-length Nfs1-HA_2_. Protein expression was induced by 1 mM IPTG, and bacterial pellets were lysed in PBS by sonication and clarified by centrifugation. Ni-NTA resin was used to pull-down His_6_-tagged proteins from clarified lysates, and proteins were eluted in 500 mM imidazole. To purify Isd11-containing complexes, the imidazole eluate was run over an S-100 size-exclusion column on an AKTA FPLC system. Lysates or eluates were diluted into SDS sample buffer.

Samples were fractionated by SDS-polyacrylamide gel electrophoresis (PAGE) using 10% acrylamide gels run at 120 V in the BIO-RAD mini-PROTEAN electrophoresis system and transferred to nitrocellulose. Membranes were probed with 1:1000 dilutions of Roche rat anti-HA monoclonal 3F10 antibody, Abcam goat anti-GFP polyclonal antibody ab5450, Abcam rabbit anti-Nfs1 polyclonal antibody ab229829, Novus anti *P. falciparum* Hsp60 polyclonal antibody NBP2-12734, and/or rabbit anti-EF1α or EF1β polyclonal antibody (65). Blots were probed with 1:5000 dilutions of fluorophore-conjugated secondary antibodies and imaged on a Licor Odyssey system.

### Microscopy

For live-cell experiments, parasites nuclei were visualized by staining with Hoechst 33342, and mitochondrial polarization was monitored by incubating parasites with 10 nM MitoTracker Red CMXROS for 15 minutes prior to wash-out and imaging. 40-50 total parasites for each condition in two independent experiments were scored for focal or dispersed MitoTracker signal, and cell percentages were analyzed by two-tailed unpaired t-test in Graphpad Prism. For IFA studies, the parasite mitochondrion was visualized using the Novus anti-HSP60 antibody and AlexaFluor 647-conjugated goat anti-rabbit 2° antibody (Invitrogen Life Technologies A21244), and mACP-HA_2_ was visualized with the rat anti-HA 3F10 primary antibody and FITC-conjugated donkey anti-rat 2° antibody (Invitrogen Life Technologies A18746). Images were taken on an EVOS M5000 imaging system or Zeiss 880 laser-scanning confocal microscope fitted with an Airyscan detector. Fiji/ImageJ was used to process and analyze images. All image adjustments, including contrast and brightness, were made on a linear scale.

### Sequence analysis

Sequence similarity searches of the *P. falciparum* genome for protein homologs of *E. coli* FASII proteins and known LYR proteins from yeast and humans (31) were performed by BLASTP analysis as implemented at the Plasmodium Genomics Resource webpage (www.plasmodb.org, release 48). Only protein hits with e-values <0.01 were retained.

### Structural modeling

A homology model of *P. falciparum* mACP bound to Isd11 was created using the InterPred modeling interface (66) and the previously determined structure (25) of *E. coli* ACP bound to human Isd11 (PDB 5USR, chains L and D). This structural model was independently refined using Rosetta Dual Relax (67), with and without restraining refinement to template coordinates, and evaluated for Rosetta energy (68). The unrestrained refinement resulted in the final model of lowest calculated energy, and this model was used for further analysis. MacPyMOL version 1.8 was used for structural visualization.

## ACKNOWLEDGEMENTS

We thank Greg Ducker, Chris Hill, Roland Lill, Sara Nowinski, Jared Rutter, Dennis Winge, and members of the Sigala lab for helpful discussions and thank Sean Prigge for helpful discussions and the plasmid encoding *P. falciparum* Isd11-GFP. We thank Rebecca Marvin for assistance with making the *P. falciparum* cytochrome c_1_-GFP expression plasmid, Megan Okada for assistance with graphical figure schemes and the design of integration vectors, and Sandra Osburn for assistance with recombinant protein mass spectrometry. This work was funded by a pilot grant to P.A.S. from the Utah Center for Iron and Heme Disorders (funded by NIH grant U54 DK110858) and NIH grants to P.A.S. (R35 GM133764) and J.A.W. (R01 GM089778). P.A.S. holds a Career Award at the Scientific Interface from the Burroughs Wellcome Fund and a Pew Biomedical Scholarship from the Pew Charitable Trusts. J.S. was funded in part by NIH T32 GM122740. S.F. was supported by the African American Doctoral Scholars Initiative at the University of Utah. Microscopy, flow cytometry, protein mass spectrometry, and DNA synthesis and sequencing were performed using core facilities at the University of Utah. Mass spectrometry core facilities at the University of Utah were supported by an NIH instrument grant (S10 OD018210).

## Notes

### Competing Interest Statement

The authors have declared no competing interest.

### Summary of Updates

Title and abstract revised and main text contains minor revisions and several new citations.

