## Supporting Information for "Divergent Acyl Carrier Protein Decouples Mitochondrial Fe-S Cluster Biogenesis from Fatty Acid Synthesis in Malaria Parasites"

### MATERIALS AND METHODS

**Cloning.** For episomal protein expression, the genes encoding mitochondrial ACP (PF3D7_1208300), Rieske protein (PF3D7_1439400), and cytochrome $c_1$ (PF3D7_ 1462700) were PCR-amplified from Dd2 parasite cDNA using primer sets 1/2, 3/4, and 5/6 (Table S4), respectively, and cloned into pTEOE (1) at the XhoI/AvrII sites in frame with a C-terminal dual hemagglutinin ($HA_2$) tag (mACP and Rieske) or GFP tag (cytochrome $c_1$). The pTEOE vector contains the HSP86 promoter to drive episomal protein expression, encodes human DHFR as a positive selection cassette, and is co-transfected with plasmid pHTH that contains the piggyBac transposase (2) for integration into the parasite genome. Ligation-independent cloning was performed with the QuantaBio RepliQa HiFi Assembly Mix (VWR 95190-050). Cloning reaction mixes were transformed into Top10 chemically competent cells, and bacterial clones were selected for carbenicillin (Sigma C3416) resistance. Plasmid DNA was isolated using the PureLink Plasmid Miniprep system (Invitrogen K210011), and correct plasmid insert sequences were confirmed by Sanger sequencing (University of Utah DNA Sequencing Core) using vector-specific primers.

For recombinant protein expression in *E. coli*, the gene for *P. falciparum* Nfs1(PF3D7_0727200) was cloned from Dd2 parasite cDNA, and genes for mACP and truncated mACP (Δ2-50) were sub-cloned by PCR from the pTEOE plasmid (see above) using primer pairs 7/8, 9/10, and 11/12, respectively (Table S4). These genes were inserted into the NcoI/XhoI sites of pET28a (Novagen 69864) with a C-terminal $HA_2$ tag using ligation-independent cloning with the QuantaBio system. *P. falciparum* Isd11 was PCR-amplified from Dd2 parasite cDNA using primer sets 13/14 and cloned into the NcoI/XhoI sites of pET21d (Novagen/MilliporeSigma 69743) in-frame with the C-terminal $His_6$ tag using ligation-independent methods. Correct insert sequences were verified by Sanger sequencing of purified plasmid DNA.

**CRISPR-Cas9 genome editing:** CRISPR/Cas9-stimulated repair by double-crossover homologous recombination was used to tag the mACP gene to encode a C-terminal HA-FLAG epitope fusion tag and the 3' 10X aptamer/TetR-DOZI system (3) to enable regulated mACP expression using anhydrotetracycline (aTc, Caymen Chemicals 10009542). A guide RNA sequence corresponding to TGGTATTGTTATATTAAATT was cloned with ligation-independent methods using primer pair 15/16 (Table S4) into a modified version of the previously-published pAIO CRISPR/Cas9 vector (4) in which the BtgZI site was replaced with a unique HindIII site to facilitate cloning. To tag the mACP gene, the donor pMG75 repair plasmid was created by using ligation-independent cloning to insert a gBlock gene fragment ordered from IDT into the unique AscI/AatII cloning sites. This gBlock contained 200 bp of the 3' untranslated region of the mACP gene (starting at position 135 downstream from the TAA stop codon), an AfeI site, and the 312 bp of the 3' end of the mACP coding sequence (excluding the 129 bp intron). The gBlock sequence included a shield mutation to ablate the CRISPR PAM sequence CGG that immediately follows the gRNA sequence above in the antisense strand of the coding sequence by mutating it to CTG, resulting in a silent mutation of the Ser125 codon from TCC to TCT. Before transfection, the pMG75 vector was linearized by AfeI digestion performed overnight at 37° C, followed by deactivation with Antarctic Phosphatase (NEB M0289S).

**Site-directed mutagenesis:** The mACP Phe113Ala (pTEOE) and Isd11 LYR12-14AAA (pET21d) mutations were introduced by PCR using primer pairs 17/18 and 19/20 (Table S4), respectively. For mACP, primer pairs 9/18 and 17/10 were used to introduce the Phe113Ala mutation and generate two insert fragments by PCR using the mACP-pET28a plasmid as template. The two fragments were joined by overlap sewing PCR using primer set 9/10. The same PCR process and primers pairs 13/20 and 19/14 were used to make Isd11 LYR12-14AAA using Isd11-pET21d as the template. Primer pairs 24/14, 25/14, and 26/14 were used to make the respective single Isd11 LYR mutations of L12A, Y13A, and R14A. Correct insert sequences for the mutated genes were confirmed by Sanger sequencing of plasmid DNA.

**Parasite culturing and transfection:** All experiments were performed using *Plasmodium falciparum* Dd2 parasites (5), whose identity was confirmed based on expected drug resistance. Parasite cultures were mycoplasma-free by PCR test. Parasite culturing was performed in Roswell Park Memorial Institute medium (RPMI-1640, Thermo Fisher 23400021) supplemented with 2.5 g/L Albumax I Lipid-Rich BSA (Thermo Fisher 11020039), 15 mg/L hypoxanthine (Sigma H9636), 110 mg/L sodium pyruvate (Sigma P5280), 1.19 g/L HEPES (Sigma H4034), 2.52 g/L sodium bicarbonate (Sigma S5761), 2 g/L glucose (Sigma G7021), and 10 mg/L gentamicin (Invitrogen Life Technologies 15750060). Cultures were maintained at 2% hematocrit in human erythrocytes obtained from the University of Utah Hospital blood bank, at 37 ˚C, and 5% $CO_2$.

For episomal protein expression using the pTEOE vector, parasite-infected erythrocytes were transfected in 1X cytomix containing 50-100 µg of purified plasmids and 25 µg of the pHTH transposase plasmid by electroporation in 0.2 cm cuvettes using a Bio-Rad Gene Pulser Xcell system (0.31 kV, 925 µF). Transfected cultures were allowed to expand in the absence of drug for

48 hours and then selected in 5 nM WR99210 (Jacobus Pharmaceuticals). Stable drug-resistant parasites returned from transfection in 2-8 weeks. The pRL2 plasmid encoding Isd11-GFP (6) was a gift from Sean Prigge (Johns Hopkins University). This plasmid was transfected into stably selected mACP-$HA_2$ (pTEOE) Dd2 parasites and selected with 5 nM WR99210 and 6 μM blasticidin-S (ThermoFisher/Gibco R21001). Similarly, plasmids for Rieske-$HA_2$, mACP-$HA_2$ and Phe113Ala mACP-$HA_2$ were separately transfected into the polyclonal Dd2 mACP-HA-FLAG aptamer/TetR-DOZI knockdown line (described below) and selected with 5 nM WR99210, 6 μM blasticidin-S, and 0.5 μM aTc.

For CRISPR/Cas9-based editing of the mACP genomic locus, 50-100 μg each of the linearized pMG75 donor plasmid and pAIO plasmid (expressing gRNA-Cas9) were mixed and transfected into Dd2 parasites and selected with blasticidin-S in the presence of 0.5-1 μM aTc. Polyclonal parasites returning from transfection were genotyped by PCR using the primer sets 21/22 and 1/22 (Table S4) for the wild-type mACP gene. Primer sets 21/23 and 1/23 were used to test parasites for integration and tagging of the mACP gene (Fig. S3). This analysis indicated that DNA recombination with the 5' homology arm occurred upstream of the single intron in the WT genomic locus, resulting in the edited gene lacking this intron. A PCR amplicon for the edited but not WT (unmodified) mACP gene was detected in polyclonal parasites, which were used for most subsequent experiments. Clonal parasite integrants were isolated by limiting dilution of the polyclonal culture and were PCR-genotyped using the same primer sets as above. All growth assays involving conditional mACP expression were performed with polyclonal parasites except for decyl-ubiquinone rescue experiments in Fig. 5, which were performed with clone B5.

**Parasite growth assays:** Parasites were synchronized to rings by treatment with 5% D-sorbitol. For growth assays involving regulated mACP expression, aptamer-tagged parasites were washed 3 times after synchronization with RPMI media lacking aTc and then divided into 2 equal parts before supplementing one part with 0.5 μM aTc. Growth was monitored by diluting sorbitol-synchronized parasites to ~0.5% parasitemia and allowing culture expansion over several days with daily media changes. For decyl-ubiquinone (dQ, Caymen Chemicals 55486005) rescue experiments, dQ was dissolved in DMSO and added directly to cultures with a final concentration of 15 μM and ≤0.3% DMSO. Proguanil (Sigma 637321) was added to cultures at a final concentration of 1 μM at the beginning of growth experiments (Fig. 5B) and added at 5 μM at the end of the 1st cycle (24 hours after synchronization) in mitochondrial depolarization studies (Fig. 5C). Atovaquone (Caymen Chemicals 95233184) was used as a positive control for dQ-rescue and membrane-polarization experiments and was dissolved in DMSO and added to corresponding cultures at a final concentration of 100 nM and ≤0.3% DMSO. Parasitemia was monitored daily by flow cytometry by diluting 10 μl of each parasite culture well into 200μl of 1.0 μg/ml acridine orange (Invitrogen Life Technologies A3568) in phosphate buffered saline (PBS) and analysis on a BD FACSCelesta system monitoring SSC-A, FSC-A, PE-A, FITC-A, and PerCP-Cy5-5-A channels. Parasitemia was determined by flow cytometry in ≥2 biological replicates (distinct parasite samples set up in parallel) and reported as an average and standard deviation, as indicated in each figure legend. Graphs were plotted using GraphPad Prism 8.0.

**Immunoprecipitation experiments:** Dd2 parasites expressing mACP-$HA_2$ or endogenously tagged mACP-HA/FLAG were harvested by centrifugation, treated with 0.05% saponin (Sigma 84510) in PBS for 5 min at room temperature to lyse erythrocytes, and spun down by centrifugation at 4,000 rpm for 30 minutes at 4°C. For HA-tagged proteins, immunoprecipitation (IP) was performed using Pierce anti-HA magnetic beads (Thermo Scientific 88836). Parasite pellets from ~50 mL cultures were lysed in 1 mL 1% triton (Sigma 9002931) or 1% digitonin in cold 1X PBS plus protease inhibitor (Thermo Scientific A32955). Pellets were dispersed by brief sonication on a Branson sonicator equipped with a microtip probe and then incubated at 4°C for 1 hour on a rotator. Lysates were clarified by centrifugation at 13,000 RPM for 10 minutes. 30 μL of anti-HA magnetic beads was equilibrated in 170 μL cold 1X Tris-buffered saline (20 mM Tris, pH 7.6, 150 mM NaCl) + 0.05% Tween-20 (TBS-T), and beads were collected against a magnetic stand. Parasite lysates supernatants (1 mL) were added to the equilibrated beads and incubated for 1 hour at 4°C by rotation. Beads were washed three times with ice-cold 1X TBS-T. Bound proteins were eluted with ~100 μL 8M urea (in 100 mM Tris at pH 8.8) and stored at -20°C until use.

Parasite samples expressing GFP-tagged proteins (Isd11-GFP and cyt $c_1$-GFP) were harvested and lysed as above. IP of GFP-tagged proteins was performed using 50 μL of protein A or G Dynabeads (Thermo Scientific 10001D or 10003D) that were equilibrated as described above. 150 μL of cold 1X TBS-T plus 0.5 μL of Goat anti-GFP primary antibody (Abcam ab5450) was added to the equilibrated beads and incubated for 10 minutes before adding the clarified lysate as described above. Washes and elutions were performed as described above.

For mass spectrometry experiments, IP eluates were precipitated by adding 100% trichloroacetic acid (Sigma 76039) to a final concentration of 20% and incubated on ice for 1 hour in 1.6 mL eppendurf tubes. Tubes were spun at 13,000 RPM for 25 minutes at 4°C. Supernatants were removed by vacuum aspiration, and protein pellets were washed once with 500 μL of cold acetone. The protein pellets were air-dried for 30 minutes and stored at -20°C.

**Western blot analyses:** Samples were fractionated by SDS-polyacrylamide gel electrophoresis (PAGE) using 10% acrylamide gels run at 120 V in the BIO-RAD mini-PROTEAN electrophoresis system. For SDS-PAGE analysis of whole parasite extracts, 1X sample buffer containing beta-mercaptoethanol was added to parasite samples before heating at 95°C for 10 minutes and centrifuging at 13,000 rpm for 5 minutes. Fractionated proteins were transferred from polyacrylamide gel to nitrocellulose membrane at 100V for one hour using the BIO-RAD wet-transfer system. Membranes were blocked in 1% casein/PBS for one hour at room temperature and then probed with primary antibody overnight at 4°C and secondary antibody at room temperature for 1 hour. Samples containing HA-tagged proteins were probed with a 1:1000 dilution of Roche Rat anti-HA monoclonal 3F10 primary (Sigma 11867423001) and a 1:5000 dilution of Donkey anti-Rat DyLight800 (Invitrogen Life Technologies SA5-10032). Membranes containing GFP-tagged proteins were probed with a 1:1000 of Goat anti-GFP polyclonal antibody (Abcam ab5450) and a 1:5000 dilution of Donkey anti-Rabbit DyLight680 (Invitrogen SA5-10042). Membranes were probed for *P. falciparum* Nfs1 with a 1:1000 dilution of Rabbit anti-Nfs1 (Abcam ab229829) 1:5000 dilution of Donkey anti-Rabbit DyLight800 (Invitrogen SA5-10044). Rabbit antibodies that recognize *P. falciparum* Hsp60 (Novus NBP2-12734) or elongation factor alpha or beta (EF1α or EF1β) (7) were used as loading controls at 1:1000 dilution. Membranes were imaged using the Licor Odyssey system. All image adjustments, including contrast and brightness, were linear.

**Mass spectrometry of parasite IP samples:** For identification of mACP-interacting proteins in IP experiments, protein samples were reduced and alkylated using 5 mM Tris (2-carboxyethyl) phosphine and 10 mM iodoacetamide, respectively, and then enzymatically digested by sequential addition of trypsin and lys-C proteases, as previously described (8, 9). The digested peptides were desalted using Pierce C18 tips (Thermo Fisher Scientific), dried, and resuspended in 5% formic acid. Approximately 1 μg of digested peptides was loaded onto a 25-cm-long, 75-μm inner diameter fused silica capillary packed in-house with bulk C18 reversed phase resin (1.9 μm, 100A pores, Dr. Maisch GmbH). The 140-min water-acetonitrile gradient was delivered using a Dionex Ultimate 3,000 ultra high-performance liquid chromatography system (Thermo Fisher Scientific) at a flow rate of 200 nl/min (Buffer A: water with 3% DMSO and 0.1% formic acid, and Buffer B: acetonitrile with 3% DMSO and 0.1% formic acid). Eluted peptides were ionized by the application of distal 2.2 kV and introduced into the Orbitrap Fusion Lumos mass spectrometer (Thermo Fisher Scientific) and analyzed by tandem mass spectrometry. Data was acquired using a Data-Dependent Acquisition method consisting of a full MS1 scan (resolution = 120,000) followed by sequential MS2 scans (resolution = 15,000) to utilize the remainder of the 3-s cycle time. Data analysis was accomplished using the Integrated Proteomics Pipeline 2 (Integrated Proteomics Applications, San Diego, CA). Data was searched against the protein database from *P. falciparum* 3D7 downloaded from UniprotKB (10,826 entries) on October 2103. Tandem mass spectrometry spectra searched using the ProLuCID algorithm followed by filtering of peptide-to-spectrum matches by DTASelect using a decoy database-estimated false discovery rate of <1%.

To analyze specific enrichment of Nfs1 in the mACP-$HA_2$ IP versus aACP-$HA_2$ IP for each matched experiment, the spectral counts observed for each protein in the mACP IP were divided by the spectral counts observed for the same protein in the aACP IP. For purposes of this analysis, all zero values for spectral counts were arbitrarily converted to a one. The $\log_2$ value of the mACP/aACP spectral count ratio for each protein was plotted for each of the 3 matched experiments. Nfs1 was highly enriched in the mACP data set and among the top 0.5-1% of proteins based on spectral count ratio.

**Mass spectrometry of purified recombinant protein samples:** For experiments involving Ni-NTA pull-down of recombinant *P. falciparum* Isd1l-His$_6$ from *E. coli* co-expressing *P. falciparum* (Δ2-50) mACP (described below), purified proteins were identified by proteolytic digestion and tandem mass spectrometry. Proteins were reduced with DTT for 45 minutes at 60 °C and then alkylated with iodoacetamide for 30 minutes at room temperature. Proteins were digested overnight at 38 °C with Trypsin/LysC mixture using one µg of trypsin per sample and quenched by acidification with 1% formic acid to a pH of 2-3. Reversed-phase nano-LC/MS/MS was performed on an UltiMate 3000 RSLCnano system (Dionex) coupled to a ThermoScientific QExactive-HF mass spectrometer equipped with a nanoelectrospray source. Concentrated samples were diluted with a 1:1 ratio of sample:0.1% formic acid in water. Five µL of the samples were injected onto the liquid chromatograph. A gradient of reversed-phase buffers (Buffer A: 0.2% formic acid in water; Buffer B: 0.2% formic acid in acetonitrile) at a flow rate of 150 µL/min at 60 °C was set-up. The LC run lasted for 83 minutes with a starting concentration of 5% buffer B increasing to 55% over the initial 53 minutes and a further increase in concentration to 95% over 63 minutes. A 40 cm long/100 µm inner diameter nanocolumn was employed for chromatographic separation. The column is a reverse-phase BEH C18 3.0 µm nanocolumn. MS/MS data was acquired using an auto-MS/MS method selecting the most abundant precursor ions for fragmentation. The mass-to-charge range was set to 350-1800. Mascot generic format (MGF) files were generated from the raw MS/MS data. Mascot (version 2.6) uses the MGF file for database searching and protein identification. For these samples the Custom database was searched with the

*Plasmodium* taxonomy selected. The parameters used for the Mascot searches were: trypsin digest; two missed cleavages; carbamidomethylation of cysteine set as fixed modification; oxidation of methionine and acetylation of the n-terminus were set as variable modifications; and the maximum allowed mass deviation was set at 11 ppm.

**Sequence homology searches:** Sequence similarity searches of the *P. falciparum* genome for protein homologs of known LYR proteins from yeast and humans (10) were performed by BLASTP analysis as implemented at the Plasmodium Genomics Resource webpage (www.plasmodb.org, release 48). The amino acid sequence of each human LYR protein homolog was used as bait, based on the following Uniprot accession codes: LYRM1 (043325), LYRM2 (Q9NU23), LYRM3 (Q9Y6M9), LYRM4 (Q9HD34), LYRM5 (Q6IPRI), LYRM6 (P56556), LYRM7 (Q5U5X0), LYRM8 (A6NFY7), LYRM9 (A8MSI8), ACN9 (Q9NRP4), C7orf55 (Q96HJ9), and L0R8F8. Only protein hits with e-values <0.01 were retained.

**Fluorescence microscopy:** For live-cell experiments, parasites nuclei were visualized by incubating samples with 1-2 μg/ml Hoechst 33342 (Thermo Scientific Pierce 62249) for 10-20 minutes at room temperature. The parasite mitochondrion was visualized by incubating parasites with 10 nM MitoTracker Red CMXROS (Invitrogen Life Technologies M7512) for 15 minutes prior to wash-out and imaging. 40-50 total parasites for each condition in two independent experiments were scored for focal or dispersed MitoTracker signal, and cell percentages were analyzed by two-tailed unpaired t-test in Graphpad Prism. For immunofluorescence assay (IFA) experiments, parasites were fixed, stained, and mounted, as previously described (1, 11). For IFA studies, the parasite mitochondrion was visualized using a polyclonal Rabbit anti-HSP60 antibody (Novus NBP2-12734) and AlexaFluor 647-conjugated Goat anti-Rabbit 2˚ antibody (Invitrogen Life Technologies A21244), the nucleus was stained with ProLong Gold Antifade Mountant with DAPI (Invitrogen Life Technologies P36931), and mACP-$HA_2$ was visualized with a Roche Rat anti-HA monoclonal 3F10 primary antibody and FITC-conjugated Donkey anti-Rat 2˚ antibody (Invitrogen Life Technologies A18746). Images were taken on DIC/brightfield, DAPI, GFP, and RFP channels using an EVOS M5000 imaging system or Zeiss 880 Laser-Scanning Confocal Microscope fitted with an Airyscan detector. Fiji/ImageJ was used to process and analyze images. All image adjustments, including contrast and brightness, were made on a linear scale.

**Rosetta modeling of the mACP-Isd11 interface:** A homology model of *P. falciparum* mACP bound to Isd11 was created using the InterPred modeling interface (12) and the structure of bovine Complex I (13) that contained mACP bound to mammalian LYR proteins NDUFA6 and NDUFB9 (PDB 5LDW, chains T and W) as template. The homology models for ACP and Isd11 were then superimposed on the previously determined structure (14) of *E. coli* ACP bound to human Isd11 (PDB 5USR, chains L and D). The structural model for the parasite mACP-Isd11 complex was refined using Rosetta Dual Relax (15), which repeatedly alternates between coordinate minimization and side-chain packing twenty times with gradually increasing van der Waals repulsion strength. Four different modeling strategies were investigated that varied the template chains in 5USR (L and D or J and H) and whether or not to restrain the refinement on the template coordinates. Ten independent refinements were performed for each modeling strategy and evaluated for Rosetta energy (16). The unrestrained refinement of the starting model using chains

L and D resulted in the final model of lowest calculated energy, and this model was used for further analysis. MacPyMOL (Schrodinger) version 1.8 was used for structural visualization.

**Recombinant protein expression in *E. coli* and purification:** Chemically competent *E. coli* BL21/DE3 cells were transformed with mACP-$HA_2$/pET28a (full-length or the Δ2-50 truncation and WT or F113A mutant), with or without Isd11-$His_6$/pET21d (WT, YR/AA, or LYR/AAA mutants), and with or without full-length Nfs1-$HA_2$/pET28a plasmids by heat shock. The bacteria were grown at 37°C in 20-mL LB media in the presence of ampicillin (100 μg/mL) or kanamycin (50 μg/mL). Bacterial cultures were allowed to grow to an optical density (at 600 nm) of 0.4 – 0.6 before inducing protein expression with isopropyl 1-thio-β-galactopyranoside (IPTG) (Goldbio 367931) at a final concentration of 1 mM. Induced cultures were grown overnight at 20°C before harvesting by centrifugation. Bacterial pellets were either used immediately or stored at -20°C. Bacterial pellets were resuspended in 1 mL 1X cold PBS and lysed by sonication on ice using a Branson sonicator equipped with a microtip for 5 sets of 10 pulses at 50% power and 50% duty cycle. Supernatants were clarified by centrifugation at 13,000 RPM for 10 minutes. For purifying hexa-His-tagged proteins, 100 μL of Ni-NTA resin (Thermo Scientific 88221) was equilibrated with 1 mL of buffer A (50 mM $NaH_2PO_4.H_20$, 500 mM NaCl, and 5 mM imidazole, pH 8.0). 100 μL of the clarified lysate supernatant was added to the equilibrated resin, diluted to 1 mL with buffer A, and incubated for 30 minutes. Resin was collected by centrifugation and washed three times with buffer A and once with wash buffer (Buffer A plus 50 mM imidazole). Bound protein was eluted by incubating resin with 100 μL elution buffer B (Buffer A plus 500 mM imidazole) for 15 minutes on ice. To purify Isd11-containing complexes in lysates from bacteria co-expressing *P. falciparum* $His_6$-Isd11 and Δ2-50 mACP, the imidazole eluate was run over an S-100 size-exclusion column (Cytiva Life Sciences 17116501) on an AKTA FPLC system (Cytiva Life Sciences). Lysates or eluates were diluted into SDS sample buffer and analyzed by SDS-PAGE and western blot analysis.

**Anti-Nfs1 antibody validation:** The commercial Rabbit anti-Nfs1 primary antibody (Abcam ab229829) was reported to be raised against the amino acids 208-457 of human Nfs1, which is 55% identical to *P. falciparum* Nfs1 when aligned by BLAST. To determine if this commercial antibody selectively recognized *P. falciparum* Nfs1, we recombinantly expressed full-length *P. falciparum* Nfs1 or mACP in *E. coli* using the Nfs1-pET28a or mACP-pET28a constructs with a C-terminal dual $HA_2$ tag (described above) that results in protein expression with expected sizes of 65 kDa (Nfs1) or 21 kDa (mACP). Protein expression was induced with IPTG, and bacteria were harvested by centrifugation and lysed in PBS by sonication. Bacterial lysate supernatants were fractionated by 10% SDS-PAGE by diluting 3 μL of clarified lysate into 47 μL of 1X SDS sample buffer, denaturing at 95°C for 10 min, spinning at 13,000 rpm for 5 min to clarify, and loading samples onto a 10% SDS-PAGE gel. After electrophoresis, samples were transferred to nitrocellulose membrane and probed with 1:1000 dilution of Roche Rat anti-HA primary antibody/1:10,000 dilution of Goat anti-Rat IRDye 680LT secondary antibody (LiCor 92668029) and 1:1000 dilution of Rabbit anti-Nfs1 primary antibody /1:10,000 dilution of Donkey anti-Rabbit IRDye800CW secondary antibody (LiCor 92632213). Detection of a single band at 65 kDa by both anti-HA and anti-Nfs1 antibodies confirmed the ability of these antibodies to recognize the HA-tagged *P. falciparum* Nfs1. The anti-Nfs1 antibody did not recognize recombinant mACP-$HA_2$, confirming its specificity, but did recognize an endogenous *E. coli* protein ~48 kDa that we identified by mass spectrometry as bacterial IscS, which is 60% identical to human Nfs1.

A

```
Yeast mACP     -------------MFRSVCRISSRV-------APSAY-RTIMG-----RSVMSN------ 28
   Pf aACP     ------------MKILLLCIIFLYYVNAFKNTQKDGVSLQILKKK---RSNQVNFLNRKN 45
   Pf mACP     MRRNIIQKILFNNKNITLCNT-----------YKRGFIQVLLKHGMNSKGIQNGFYNN-- 47
                                :*                .    ::      :.   .

Yeast mACP     --TIL---AQRFYS-------ANLSKDQVSQRVIDVIKAFDKNSPNIANKQISSDTQFHK 76
   Pf aACP     DYNLIKNKNPS----------SSL-KS-----TFDDIKKIISKQLSVEEDKIQMNSNFTK 89
   Pf mACP     KYILSAQKNSAFFSTEKSQDLSSLSKEQIEEKILTVLKKYLPPDVEIKYDEELEKYNTKD 107
                  :                 :.* *.      :  :*     . .:  .:   . :  .

Yeast mACP     DLGLDSLDTVELLVAIEEEFDIEIPDKVADELRSVGETVDYIASNPDAN-  125
   Pf aACP     DLGADSLDLVELIMALEEKFNVTISDQDALKINTVQDAIDYIEKNNKQ--  137
   Pf mACP     NRAWDFLDTVEFLIDIESEFNITIPDETADNIKTVQEVIDYIIQLNIKKT  157
```

B

putative processing site

**MRRNIIQKILFNNKNITLCNTYKRGFIQVLLKHGMNSKGIQNGFYNNKYI|LSAQKNSAFF**
**STEKSQDLSSLSKEQIEEKILTVLKKYLPPDVEIKYDEELEKYNTKDNRAWDFLDTVEFL**
**IDIESEFNITIPDETADNIKTVQEVIDYIIQLNIKKT**

**Fig. S1.** Sequence and mass spectrometry (MS) analyses. (A) Alignment of yeast ACP with *P. falciparum* apicoplast ACP (aACP) and mitochondrial ACP (mACP). (B) Sequence of *P. falciparum* mACP with peptide residues in red that were detected by tandem MS analysis of parasite samples in this study or available on PlasmoDB. The putative mACP processing site is shown, based on its proximity to the most N-terminal tryptic peptide detected (NSAFFSTEK**)** and alignment with the known mitochondrial ACP processing sites in yeast and humans (17, 18).

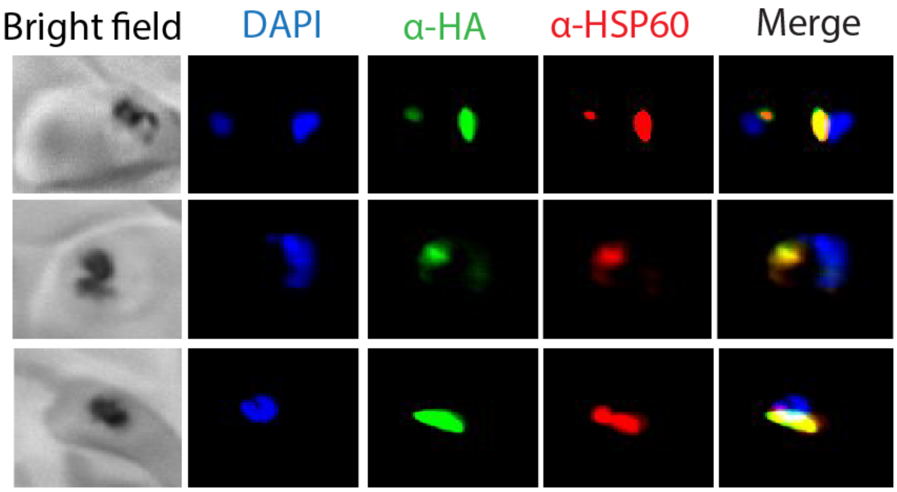


**Fig. S2.** Additional immunofluorescence microscopy images of fixed Dd2 parasites episomally expressing mACP-$HA_2$ and stained with DAPI (nucleus, blue), anti-HA (green), and anti-HSP60 (red) antibodies.

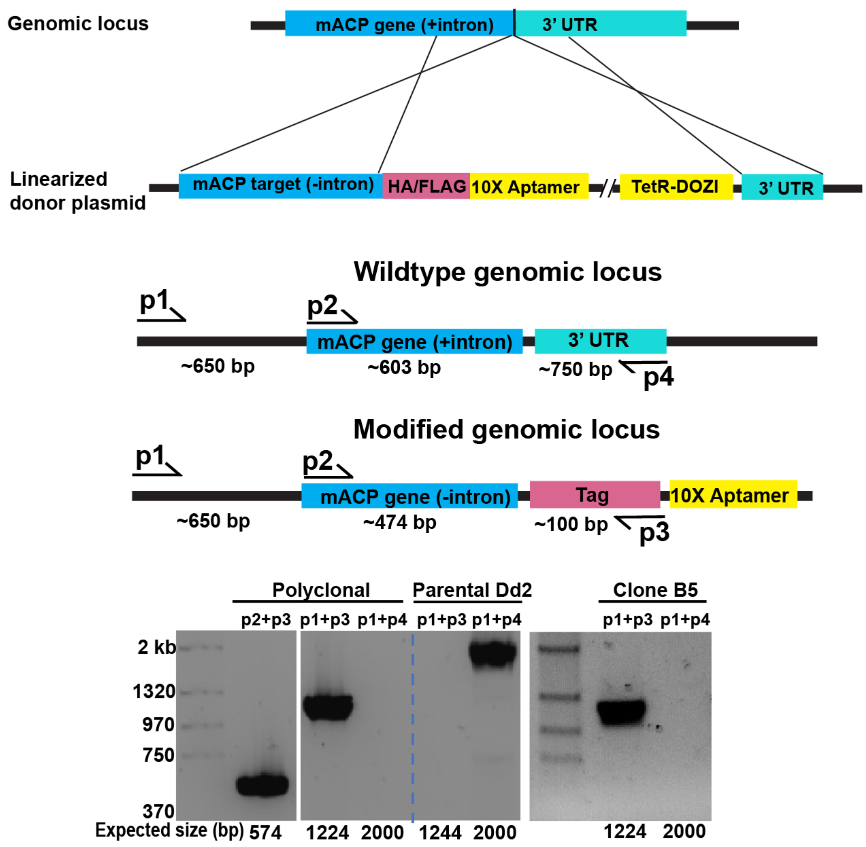


**Fig. S3.** Schematic depiction of mACP gene editing and verifying genomic integration by PCR.

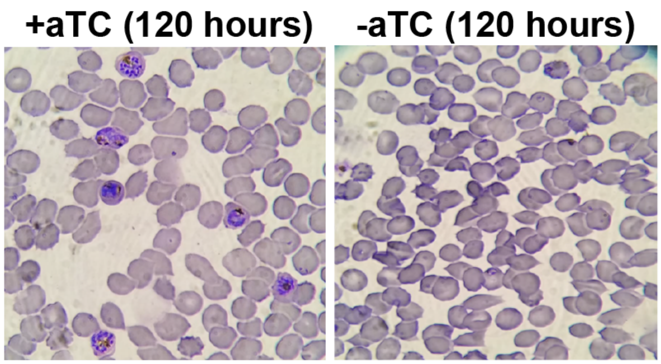


**Fig. S4.** Giemsa-stained blood smears of mACP-aptamer/TetR-DOZI parasites cultured for 120 hours (5 days) ±aTc.

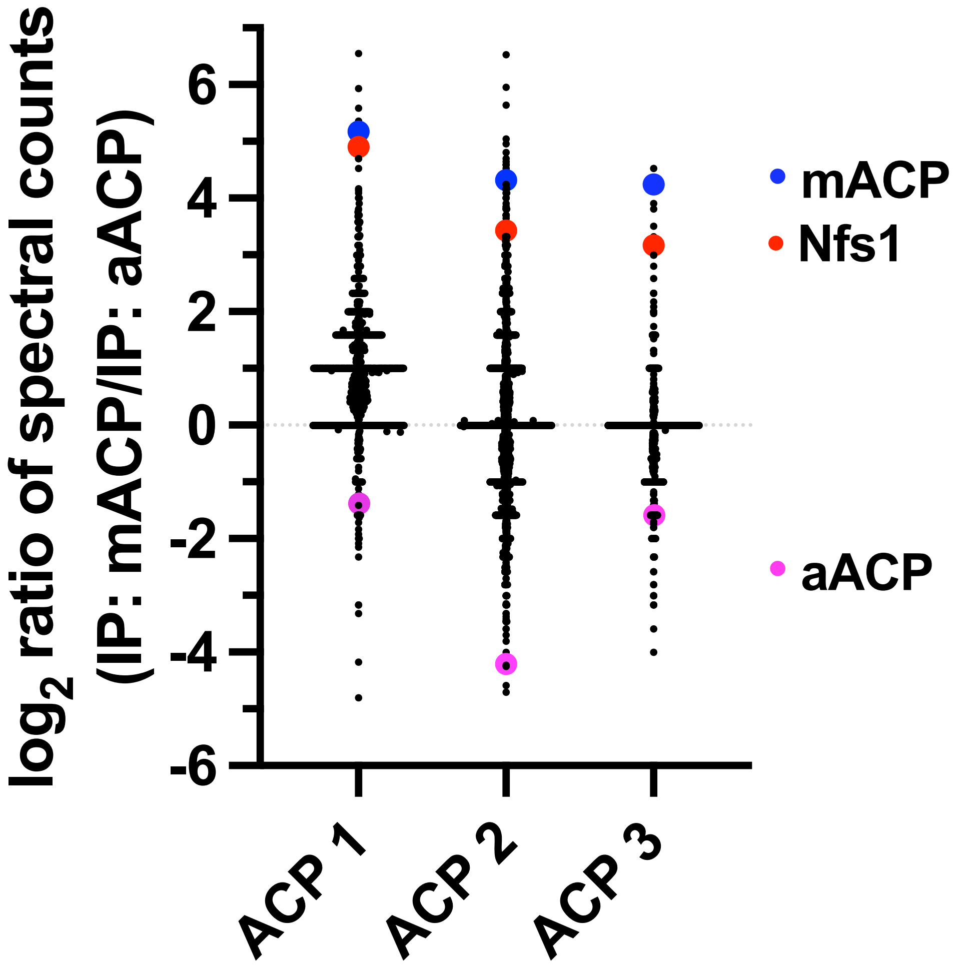


**Fig. S5.** Specific enrichment of Nfs1 for anti-HA-tag IP of mACP-$HA_2$ versus aACP-$HA_2$ in three matched experimental samples, calculated as $\log_2$ of the spectral count ratio (IP:mACP/IP: aACP).

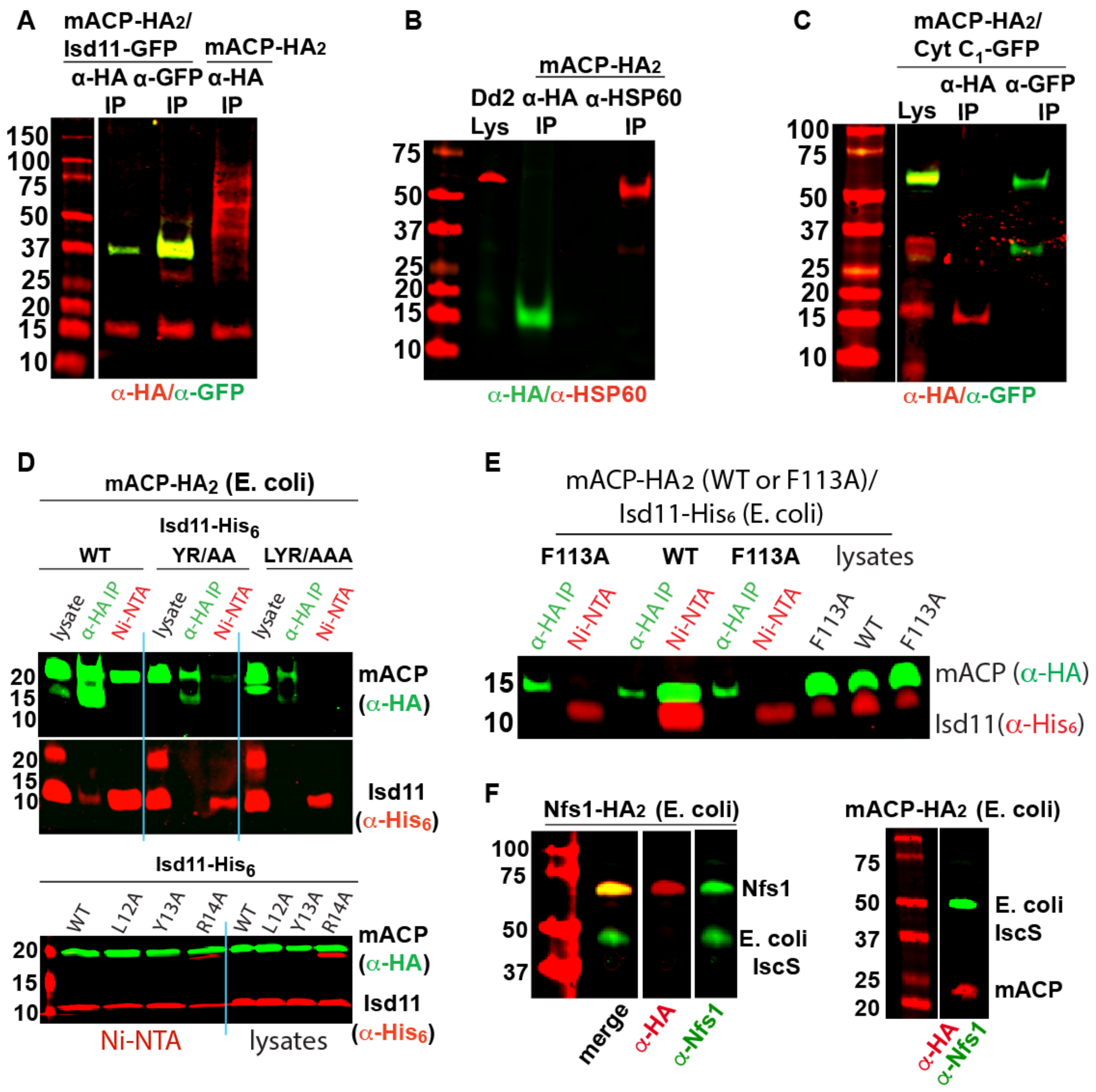


**Fig. S6.** Western blot analyses. Reciprocal immunoprecipitation/western blot studies of Dd2 parasites expressing (A) mACP-HA$_2$ with and without Isd11-GFP (B) mACP-HA$_2$ and HSP60 and (C) mACP-HA$_2$ and cyt c$_1$-GFP. Anti-HA IP or nickel-nitrilotriacetic acid (Ni-NTA) pulldown and WB studies of *E. coli* bacteria recombinantly expressing (D) full-length mACP-HA$_2$ and Isd11-His$_6$ (WT, YR/AA, LYR/AAA, L12A, Y13A, or R14A mutants) and (E) Δ2-50 mACP-HA$_2$ (WT or F113A mutant) and Isd11-His$_6$ and probed with anti-HA and anti-His$_6$ antibodies. (F) *P. falciparum* Nfs1-HA$_2$ or full-length mACP-HA$_2$. For (E) *P. falciparum* Nfs1-HA$_2$ was recognized by both the anti-HA and anti-Nfs1 antibodies whereas full-length mACP-HA$_2$ was only recognized by the anti-HA antibody. In both lysates, the anti-Nfs1 antibody also recognized the endogenous *E. coli* IscS (~48 kDa) that is ~60% identical to human Nfs1. Blots were probed as labeled and as described in Materials and Methods.

**Isd11**

MNGNQIKQLKKLYRHILNEASKFENINYNVYFSNKAKEKFREFCSDTNFESEKLKTFQNECWDYLNMLKR
QTIIHNLYHVDKPLVNK

**mACP**

LSAQKNSAFFSTEKSQDLSSLSKEQIEEKILTVLKKYLPPDVEIKYDEELEKYNTKDNRAWDFLDTVEFL
IDIESEFNITIPDETADNIKTVQEVIDYIIQLNIKKT

**Fig. S7.** Sequences of *P. falciparum* Isd11 and mACP constructs that were heterologously co-expressed *E. coli*, purified by Ni-NTA and size-exclusion chromatography, and analyzed by tandem mass spectrometry (MS). Underlined sequence reflects tryptic peptide residues that were detected by tandem MS analysis.

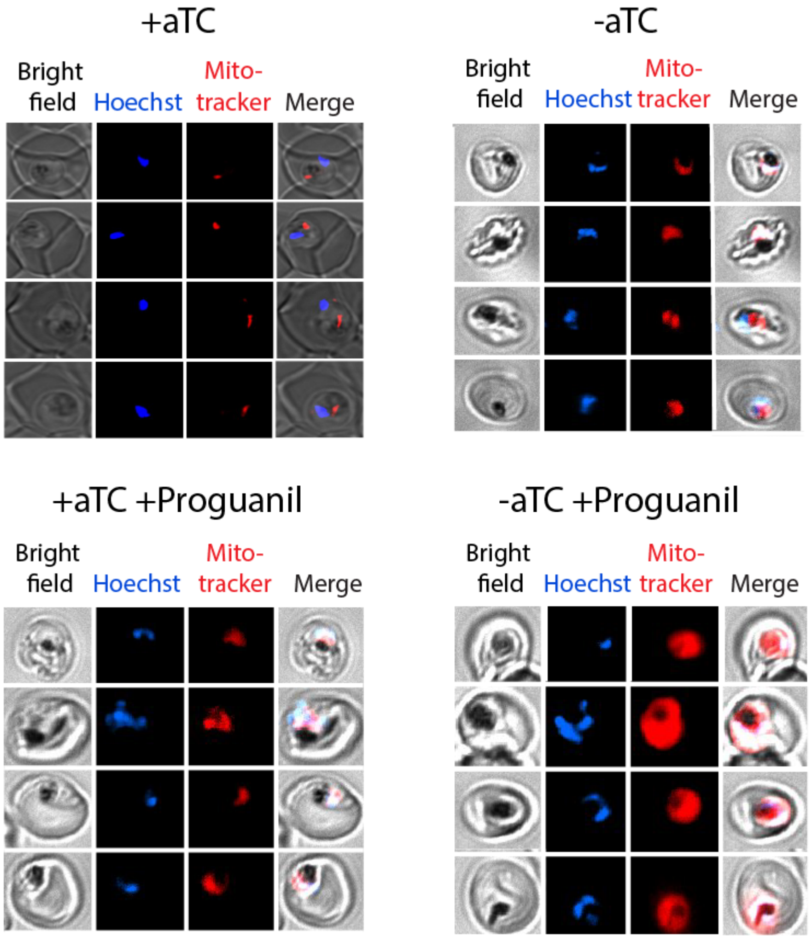


**Fig. S8.** Additional live microscopy images of mACP-aptamer/TetR-DOZI parasites cultured for 72 hours ±aTc and for 24 hours ±proguanil and treated with 10 nM Mitotracker-Red and Hoechst.

| **FASII proteins** | ***E. coli*** | ***P. falciparum* apicoplast** | **E-value** | ***P. falciparum* mitochondrion** | **E-value** |
|---|---|---|---|---|---|
| Acyl carrier protein | AcpP P0A6A8 | ACP PF3D7_0208500 | 5e-22 | ACP PF3D7_1208300 | 3e-09 |
| Phosphopantetheine transferase | AcpS P24224 | ACPS PF3D7_0420200 | 8e-15 | - | - |
| Acetyl-CoA carboxylase | AccC P24182 | ACC PF3D7_1469600 | 7e-58 | - | - |
| Malonyl-CoA:ACP transferase | FabD P0AAI9 | MCAT/FabD PF3D7_1312000 | 4e-39 | - | - |
| 3-Ketoacyl-ACP synthase | FabB/FabF P0A953 | FabB/FabF PF3D7_0626300 | 1e-48 | - | - |
| 3-Ketoacyl-ACP reductase | FabG P0AEK2 | FabG PF3D7_0922900 | 4e-80 | - | - |
| 3-Hydroxyacyl-ACP dehydratase | FabZ P0A6Q6 | FabZ PF3D7_1323000 | 3e-39 | - | - |
| 2-Enoyl-ACP reductase | FabI P0AEK4 | FabI PF3D7_0615100 | 4e-19 | - | - |
| beta-ketoacyl-ACP synthase III | FabH P0A6R0 | FabH PF3D7_0211400 | 9e-68 | - | - |

**Table S1.** *P. falciparum* homologs of *E. coli* FASII proteins.

| **LYR Protein Human/Yeast** | ***P. falciparum* homolog** | **E-value** |
|---|---|---|
| LYRM4/ISD11 | Isd11<br>PF3D7_1311000 | 3e-07/3e-06 |
| LYRM1 | - | - |
| LYRM2 | - | - |
| LYRM3 | - | - |
| LYRM5 | - | - |
| LYRM6 | - | - |
| LYRM7/MZM1 | - | - |
| LYRM8/Sdh6 | - | - |
| LYRM9 | - | - |
| ACN9/Sdh7 | - | - |
| C7orf55/FMC1 | - | - |
| L0R8F8 | - | - |

**Table S2.** BLAST analysis of LYR-motif protein homologs found in *P. falciparum*.

| | **ACP bait spectral counts** | | |
|---|---|---|---|
| | Exp. 1 | Exp. 2 | Exp. 3 |
| | Triton | Triton | Digitonin |
| **IP: mACP** | 73 | 37 | 19 |
| **IP: aACP** | 63 | 65 | 12 |

**Table S3.** Table of spectral counts for *P. falciparum* mACP or aACP detected by tandem mass spectrometry for anti-HA immunoprecipitation studies of lysates from Dd2 parasites episomally expressing mACP-$HA_2$ or aACP-$HA_2$. Parasites were lysed in either Triton X-100 or digitonin.

| No. | Primer Name | Sequence (5' to 3') |
|---|---|---|
| 1 | mACP-HA2-TEOE f | ACGATTTTTTCTCGAGATGAGAAGAAATATTATTCAAAAAATTCTGTT CAATAATA |
| 2 | mACP-HA2-TEOE r | CGTATGGGTACCTAGGTGTTTTTTTATATTTAATTGAATTATATAAT CTATAACT |
| 3 | Rieske-HA2-TEOE f | CATTTCGAATAAACACGATTTTTTCTCGAGATGAATAATATTAAATAT GTGGAACTTTTTTAC |
| 4 | Rieske-HA2-TEOE r | GTAATCTGGAACATCGTATGGGTACCTAGGTCCAATTTTTATCGTATT TTCATCAACAAATTC |
| 5 | Cyt C1-GFP-TEOE f | ACGATTTTTTCTCGAGATGGCTGGTGGGGGAG |
| 6 | Cyt C1-GFP-TEOE r | CTGCACCTGGCCTAGGTAAATATTTGATTTTTCCAAAGTCAATTCTTC TAGTTG |
| 7 | Nfs1-HA2-pET28a f | GTTTAACTTTAAGAAGGAGATATACCATGGCCAAATTTCTTCAAATA ATAAAACATCTCAAATTACAAAACAAAAAGAACGCAC |
| 8 | Nfs1-HA2-pET28a r | GATCTCAGTGGTGGTGGTGGTGGTGCTCGAGTGTCCATATAAACTTT GGAATGTCGTCATTCGAGGG |
| 9 | mACP-HA2-pET28a f | AGGAGATATACCATGGCCAGAAGAAATATTATTCAAAAAATTCTGTT CAATAATAAAAATATAACACTATGTAATACG |
| 10 | mACP-HA2-pET28a r | GGTGGTGGTGCTCGAGTTACGCGTAATCAGGTACATCGTATGGATAA GCGTAATC |
| 11 | TrmACP-HA2-pET28a f | AACTTTAAGAAGGAGATATACCATGGCCTTGAGCGCTCAAAAAAATA GTGCCTTTTTTTC |
| 12 | TrmACP-HA2-pET28a r | ATCTCAGTGGTGGTGGTGGTGGTGCTCGAGTTACGCGTAATCAGGTA CATCGTATGGATAAGC |
| 13 | Isd11-His6-pET21d f | GTTAATATTATACAATATACCTAGGATGAATGGGAATC |
| 14 | Isd11-His6-pET21d r | CAGTGAAAAGTTCTTCTCCTTTACTCGTACGTTTGTTAACTAAAGG |
| 15 | mACP_gRNA_sense | TAAGTATATAATATTTGGTATTGTTATATTAAATTGTTTTAGAGCTAG AA |
| 16 | mACP-_gRNA_antisense | TTCTAGCTCTAAAACAATTTAATATAACAATACCAAATATTATATACT TA |
| 17 | Phe-Ala mACP-HA2 f | GGATAATCGAGCCTGGGATGCTCTTGATACCGTTGAATTTTTAATAG ATATTG |
| 18 | Phe-Ala mACP-HA2 r | CAATATCTATTAAAAATTCAACGGTATCAAGAGCATCCCAGGCTCGA TTATCC |
| 19 | LYR-AAA Isd11 f | GTTAATATTATACAATATACCTAGGATGAATGGGAATCAAATAAAAC AACTTAAGAAGGCTGCCGCGCATATATTGAATGAAGC |
| 20 | LYR-AAA Isd11 r | GAAGCTTCATTCAATATATGCGCGGCAGCCTTCTTAAG |
| 21 | mACP 5'UTR f | CCATACATATTTGTGTATGTATGTATGTATTTATTTATTATGTATTCAA ATTATGC |
| 22 | mACP 3'UTR r | TTCTTCTCATCTTAAGCTATTTTATATTTATAATATAAATGAGCTCTTC AAAAGTAG |
| 23 | 3' Aptamer tag r | CCCAGGCCTCTAGTTTAC |
| 24 | L12A Isd11 f | GTTTAACTTTAAGAAGGAGATATACCATGGCCAATGGGAATCAAATA AAACAACTTAAGAAGGCTTATCGTCATATATTGAATGAAGCTTC |
| 25 | Y13A Isd11 f | GTTTAACTTTAAGAAGGAGATATACCATGGCCAATGGGAATCAAATA AAACAACTTAAGAAGTTAGCCCGTCATATATTGAATGAAGCTTC |
| 26 | R14A Isd11 f | GTTTAACTTTAAGAAGGAGATATACCATGGCCAATGGGAATCAAATA AAACAACTTAAGAAGTTATATGCACATATATTGAATGAAGCTTC |

**Table S4.** PCR primers used for cloning and genotyping.
